# Statistics of chromatin organization during cell differentiation revealed by heterogeneous cross-linked polymers

**DOI:** 10.1101/235051

**Authors:** O. Shukron, V. Piras, D Noordermeer, D. Holcman

## Abstract

Chromatin of mammalian nucleus folds into discrete contact enriched regions such as Topologically Associating domains (TADs). The folding hierarchy of the TADs and their internal organization is highly dynamic through cellular differentiation, where structural changes within and between TADs are correlated with gene activation and silencing. To account for multiple interacting TADs, we developed a parsimonious randomly cross-linked (RCL) polymer model that maps high frequency encounters present in HiC data within and between TADs into direct local monomer interactions, characterized by the number of cross-links at a given base-pair resolution. We reconstruct three TADs obtained from the mammalian X chromosome for three stages of differentiation. We compute the radius of gyration of TADs and the encounter probability between genomic segments. We found 1) a synchronous compaction and decompaction of TADs throughout differentiation and 2) secondary structures such as meta-TADs in 5C data resulting from weak inter-TAD interactions. Finally, the present method links steady-state to dynamic properties of the chromatin revealed by the distribution of anomalous exponents of single loci trajectories, reconstructed from HiC data.

## INTRODUCTION

Mammalian chromosomes fold into discrete mega-basepairs (Mbp) contact enriched regions termed Topologically associating domain (TADs). Although the precise role of TADs remain unclear, they participate in synchronous gene regulation [1, 2] and replication [3]. Gene regulation within TAD is modulated by transient loops at sub-TAD scale [1, 4–6], formed by regulatory elements such as enhancers and promoters [7]. However, sparse connectors between TADs (at a scale *>*Mbps) can significantly influence chromatin dynamics and gene regulation within TADs [8, 9]. We present and apply here methodology based on polymer model that accounts for interactions between TADs to study chromatin dynamic at long scales, an area that remains largely unexplored.

Genome organization can be probed by chromosome conformation capture (CC) methods [2, 4, 10], which report simultaneous genomic contacts (loops) at scales of kilo bps (kbps) to Mbps. TADs appear as matrix block in the contact maps by summing encounter events over an ensemble of millions of cells [1, 10]. Conformation capture contact maps provide a statistical summary of steady-state looping frequencies, but do not contain neither direct information about the size of the folded genomic section nor any transient genomic encounter times [11]. Throughout cell differentiation stages, the boundaries between TADs remain stable, [1, 12] but their internal looping pattern is highly variable. Moreover, TADs can form hierarchy structures into meta TADs, formed by inter TAD connectivity, correlated with transcription state of the chromatin [12].

Coarse-grained polymer models are used to study steady-state and transient properties of chromatin at a given scale. Starting with the Rouse polymer [13], composed of *N* monomers connected sequentially by harmonic springs, the linear connectivity in the Rouse model does not account for the complexity of molecular interactions and therefore cannot account for contact enriched regions such as TADs. To include short and long-range looping, other polymer models have been developed, with self-avoiding interactions [11, 14–22], random loops [8, 14, 23, 24] or epigenomic state [25, 26], that in addition account for TADs.

To account for interactions between multiple TADs, we develop a parsimonious polymer model of the chromatin. We extend the construction of the randomly cross-linked (RCL) polymer model, introduced in [9, 11], and account for multiple connected TADs of various sizes and connectivity within and between TADs. Random cross-links can be generated by binding molecules such as CTCF [1, 27] or by loop extrusion mechanism [28], although the exact mechanism by which they form is not really important for the present work. The random positions of the cross-links capture the heterogeneity of chromatin structure sampled in a large ensemble of cells.

We then introduce our methodology to construct an RCL polymer from the empirical encounter probability (EP) extracted from 5C or HiC data, we determine their geometrical organization and characterize their distribution in space by computing the volume of TADs using the mean radius of gyration. We further investigate the role of several parameters such as the number and positions of the cross-links or the polymer length. We show that TADs and higher-order genome organization are modulated by the inter-TAD connectivities. These connectivities determine the steady-state and time-dependent statistical properties of monomers within each TAD. We validate the present method by comparing HiC and 5C data and the reconstructed statistics between two 5C replicas. Finally, we study the reorganization of three neighboring TADs on the mammalian X chromosome throughout stages of cellular differentiation. The key step of our method was to derive a novel expression for the EP of the RCL polymer model (see Eq. 10, 12), this step was the bottleneck that we used to fit the empirical 5C data [1]. We then determine the rate of compaction and decompaction of TADs that can be correlated to gene silencing and activation, respectively. These properties cannot be studied by a direct comparison of the overall number of interactions within and between TADs from 5C data, because individual realizations cannot be discerned from the ensemble. To conclude, the present framework allows to study chromatin dynamics from HiC or 5C data, to derive its statistical properties, to connect steady-state HiC statistics with time-dependent analysis of single locus trajectories and to reveal the influence of multiple connected TADs on each other and genome reorganization throughout cellular differentiation.

## METHODS

### I. RCL POLYMER MODEL FOR MULTIPLE INTERACTING TADS

We now describe the construction of a heterogeneous Randomly Cross-Linked (RCL) polymer for multiple interacting TADs. The RCL polymer models were previously used in [11] and SI (section I) to compare the statistical properties of simulations vs. theory for a single TAD. Heterogeneous RCL polymers consist of *N*_*T*_ sequentially connected RCL polymers (Fig. 1A, SI section I) that account for connectivity within TADs of *N* = [*N*_1_, *N*_2_, …, *N*_*NT*_] monomers, respectively (Fig. 1B). The polymer is constructed as follows: the position of the ensemble of monomers ***R*** in all TADs is described by *N*_*T*_ vectors represented in a block matrix

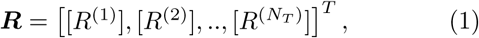

where the superscript indicates TAD 1, …, *N*_*T*_ it belongs to and square brackets indicate a block matrix, such that 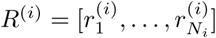. The linear backbone of the polymer is a chain of *N*_*T*_ Rouse matrices (SI Eq. 2), defined in the block matrix

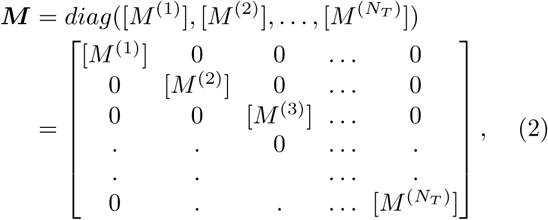

where each [*M*^(*j*)^] contains *N*_*j*_ monomers. Note, that during the numerical simulations, we connect the last monomer of block [*M*^(*j*)^] to the first one of [*M*^(*j*+1)^]. One of the key ingredients of the present model is the addition of random spring connectors between non nearest-neighbor (non-NN) monomer pairs. The maximal possible number *N*_*L*_ of non-NN connected pairs within (between) TAD *A*_*i*_ (*A*_*i*_ and *A_j_*) is given by

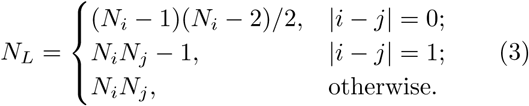

**FIG. 1.**
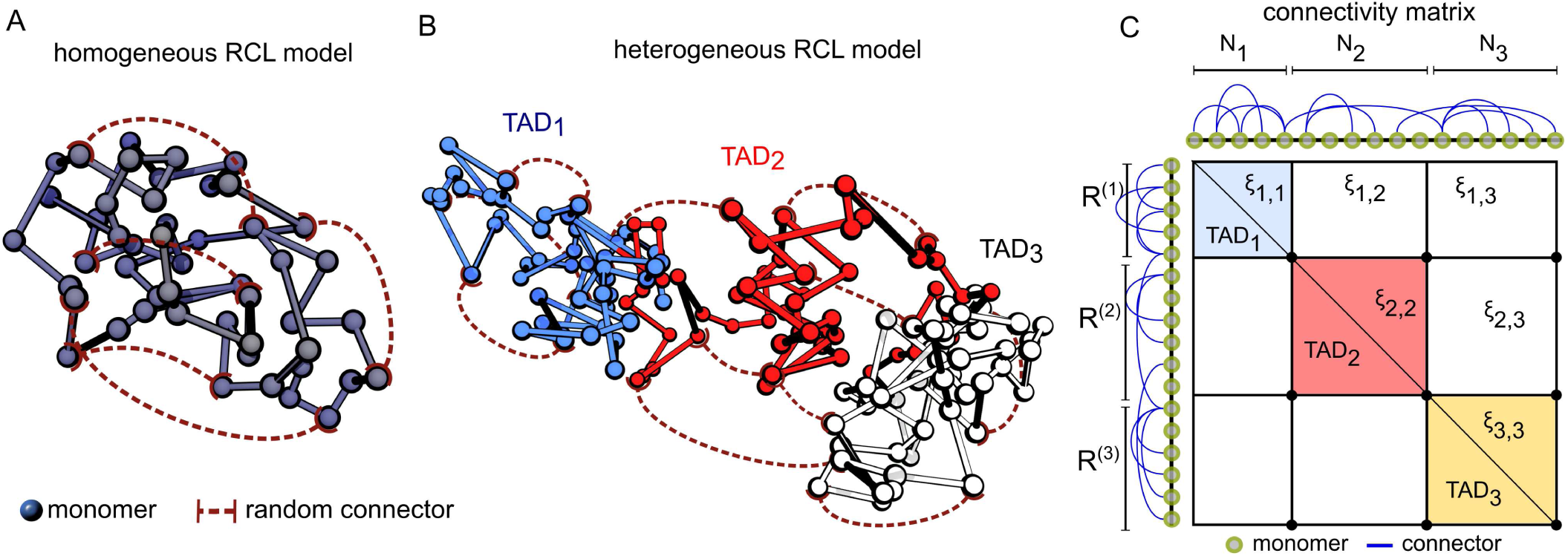
RCL polymer model for single and multiple interacting associating domains. **A.** Schematic representation of the RCL polymer for a single topologically associating domain (TAD). Monomers (circles) are connected sequentially by harmonic springs to form the polymer’s backbone (gray). Spring connectors (dashed red) are added between randomly chosen non nearest-neighbor monomers. The choice of monomer pairs to connect is randomized in each realization. All monomers share similar level of connectivity (homogeneous RCL model) **B.** Schematic representation of the RCL polymer model for three connected TADs (blue, red, white), where monomers (circles) are randomly connected (dashed red) within and between TADs by springs. Monomer connectivity within and between TADs can vary (heterogeneous RCL model). **C.** schematic representation of the symmetric connectivity matrix *B*^𝒢^(Ξ), corresponding to the connected monomers over the three TADs in panel B (diagonal bocks), where each TAD is of variable number of monomer *N* and connectivity fraction *ξ*_*ij*_.

We add *N*_*c*_(*i, j*) spring connectors randomly between non-NN monomer pairs (Fig. 1B) between TADs *A*_*i*_ *A*_*j*_. We define the square-symmetric connectivity fraction matrix Ξ = {*ξ*_*ij*_}, 1 ≤ *i, j* ≤ *N*_*T*_, as the ratio of number of connectors to the total possible number of non-NN monomer pairs

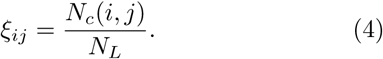

A realization 𝒢 of the RCL polymer is a random choice of *N*_*c*_(*i, j*) monomer pairs to connect. We define the added connectivity matrix *B*^𝒢^(Ξ) by

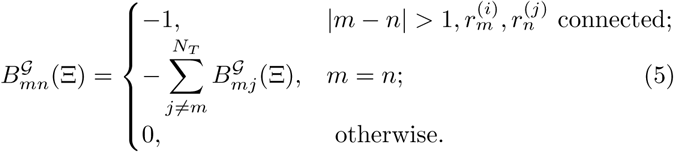

The spring constants of the linear backbone and the added *N*_*c*_ connectors are similar. The added springs keep distal connected monomers into close proximity. The energy 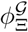 of realization 𝒢 of the RCL polymer is the sum of the spring potential of the linear backbone plus that of added random connectors:

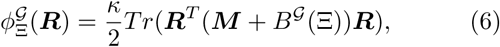

where *κ* is the spring constant, *Tr* is the trace operator. The dynamics of monomers ***R*** is induced by the potential energy 6 plus thermal fluctuations:

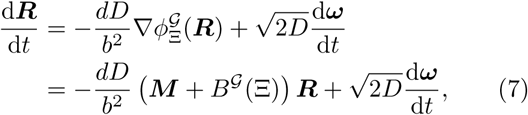

for a dimension *d* = 3, the mean distance *b* between neighboring monomers is defined for a linear chain when there are no added connectors (*Nc*=0), a diffusion coefficient *D*, and the standard *d*-dimensional Brownian motions ***ω*** have mean 0 and variance 1. We note that added connectors describe the cross-linking by binding molecules such as cohesin or CTCF, and their random positions in 𝒢 realization capture the heterogeneity over cell population. In addition, after connectors have been added the mean distance between two consecutive monomers *r*_*m*_, *r*_*m*+1_ can be smaller than the initial mean distance *b* for a Rouse chain (*Nc* = 0), as computed below, based on Eqs. 11–13.

## RESULTS

### A. Statistical properties of heterogeneous RCL and numerical validations

We will study the statistical properties of the heterogeneous RCL polymer, representing multiple TADs using first the Mean-Square-Radius of Gyration (MSRG), second the encounter probability (EP) in two cases: between monomers of the same TAD (intra TAD) and across TADs (inter-TAD), third, Mean-Square-Displacement (MSD) of single monomers, and forth, the distribution of distance between any two monomers. Details are given in the SI sections II C-II F.

1. The MSRG 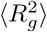 characterizes the folding of a TAD inside a ball of radius 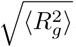. When the condition of a dominant intra-TAD connectivity (assumption *H*_1_, SI Eq. 29) is satisfied, the MSRG for TAD_*i*_ (see derivation in SI Eq. 24-34) is given by

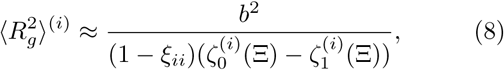

where

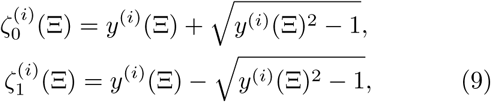

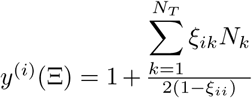, and ξ_*ij*_ is the connectivity matrix defined in 4.

2. Under the condition of non-vanishing connectivities (SI condition 50), the EP between monomer *m* and *n* within TAD *A*_*i*_ (see derivation in SI section II D) is given by

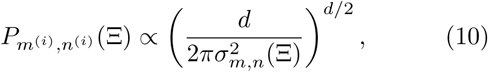

where

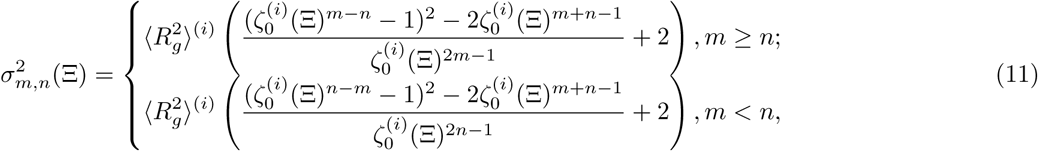

and 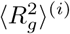 is the MSRG of TAD *A_i_*, defined by relation 8. When monomers belong to distinct TADs *i* and *j*(*i* ≠ *j*), the EP formula is modified to (see SI subsection II D)

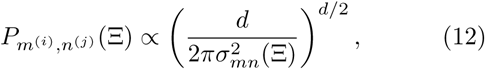

where

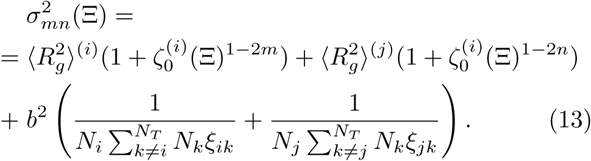

3. The MSD of a monomer 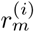 located inside *A*_*i*_ for intermediate times (see SI section II E) is given by

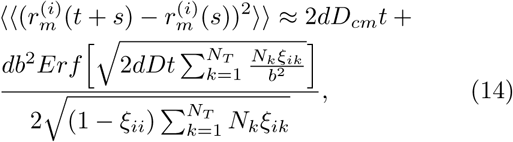

where 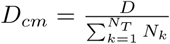 and *Erf* is the Gauss error function.

4. The distribution *f*_*D*_*mn*__(*x*) of the distance *D*_*mn*_ = ||*r*_*m*_ − *r*_*n*_|| between any two monomers *r*_*m*_ and *r*_*n*_ (see SI subsection II F) is given by

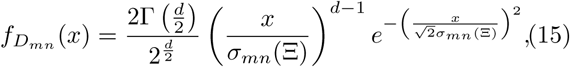

where is Γ the Γ–function.

To validate formulas 8-14 so that we can use them to extract statistical properties of 5C/HiC data, we decided to test them against numerical simulations of three synthetic interacting TADs. We constructed a RCL polymer containing three TADs with *N*_1_ = 50, *N*_2_ = 40, *N*_3_ = 60 total monomers, so that condition *H*_1_ (SI Eq. 29) about dominant intra-connectivity is satisfied. We impose the number of connectors in each TAD to be at least twice compared to the one between TADs (see Fig. 2A).

**FIG. 2.**
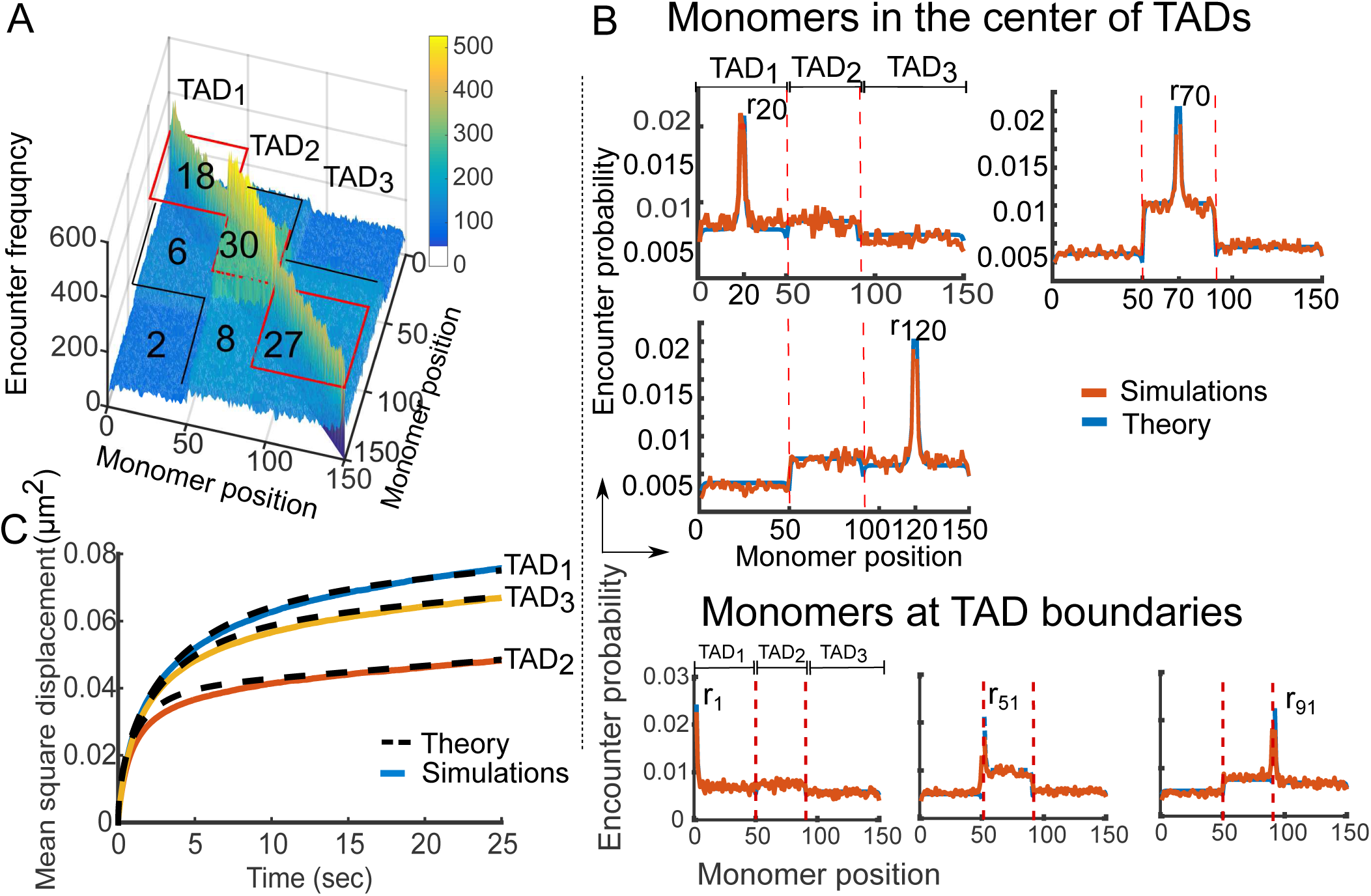
Statistical properties of the heterogeneous RCL polymer. **A.** Encounter frequency matrix of a polymer with three TAD blocks (TAD_1_, TAD_2_, TAD_3_) of *N*_1_ = 50, *N*_2_ = 40, *N*_3_ = 60 monomers, computed from 10,000 simulations of the Eq. 7 with ∆*t* = 0.01*s, D* = 8 × 10^*−*3^*µm*^2^/*s*, *d* = 3, *b* = 0.2*µm* [29]. The number of added connectors appears in each block. Three distinct diagonal TADs are visible (red boxes) where secondary structure appears (black lines) due to weak inter-TAD connectivity. **B.** Encounter probability (EP) of the heterogeneous RCL described in panel A, where the simulated (orange) and theoretical (blue, Eqs. 10, 12) EP agree. In the upper panel we plotted the EP for middle monomers in each TAD: monomer *r*_20_ (upper left), monomer *r*_70_ (top right), and monomer *r*_120_ (bottom left). In the bottom panel, we plotted the simulated (orange) vs. theoretical EPs (blue) for monomers *r*_1_ (left), *r*_51_(center), and *r*_91_(right), located at TAD boundaries. **C.** Average mean-square displacement of monomers in each TAD of the heterogeneous RCL polymer, simulated as described in panel A, simulations (continuous) vs theory (dashed, Eq. 14) until *t* = 25s.

To construct the encounter frequency matrix, we simulated equation 7 in dimension *d* = 3, with *b* = 0.2*µm* and diffusion coefficient *D* = 8× 10^−3^ *µm*^2^/*s*[29] starting with a random walk initial polymer configuration. Connectors were placed between monomers with a uniform probability in each TAD_*i*_ and in between TADs, as indicated in Fig. 2A. We ran 10,000 simulations until polymer relaxation time (see SI eq. 23 and [11]). The longest relaxation time of RCL chains containing *N*_*T*_ TADs is defined by tens of thousands of simulation steps. At the end of each realization, we collected the monomer encounters falling below the distance *ε* = 40 *nm*, and constructed the simulation encounter frequency matrix. This matrix shows three distinct diagonal blocks (Fig. 2A) resulting from high intra TAD connectivity, and further reveals a high-order organization (cyan blue in Fig. 2A), which resembles the meta-TADs discussed in [12]. We thus propose here that hierarchical TAD organization is a consequence of weak inter-connectivity properties.

We then computed the steady-state EP from the simulation encounter frequency matrix (Fig. 2A) by dividing each row with its sum. We then compared simulations and theoretical EPs (Eqs. 10 and 12) in Fig. 2B: the three sample curves for monomer *r*_20_ (upper left), *r*_70_ (upper right), and *r*_120_ (lower) located in the middle of each TAD, are in good agreement with the theory. Furthermore, the theoretical and simulated EPs for monomer *r*_1_, *r*_51_ and *r*_91_, located at boundaries of TADs (Fig. 2B bottom) are in good agreement. Finally, we computed the Mean Radius of Gyration (MRG) 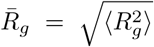, for TAD_1_, TAD_2_, and TAD_3_ given by 0.177, 0.13, 0.165*μm* (simulations), compared to 0.178, 0.13, 0.167, respectively, obtained from expression 8, which agree.

To validate the MSD expression (Eq. 14), we simulated Eq. 7 for 2500 steps with a time step Δ*t* = 0.01s, past the relaxation time *τ*(Ξ) (SI Eq. 23) and computed the average MSD over all monomers in each TAD_*i*_, *i* = 1, 2, 3. In Fig. 2C, we plotted the average MSD in each TAD against expression Eq. 14 (dashed), which are in good agreement. The overshoot of the MSD of TAD_1_, results from the weak coupling of centers of masses of TADs (see Eq. 22). The amplitude of the MSD curve is inversely proportional to the total connectivity of each TAD as shown in Fig. 2C, TAD_1_ (blue, 26 connectors), TAD_2_ (red, 44 connectors), and TAD_3_ (yellow, 37 connectors). We conclude, that the present approach (numerical and theoretical) capture the steady-state properties (Eqs. 8,10–12, 14) of multi-TAD.

In addition, we found that adding an exclusion forces with a radius of 40*nm* did not lead to any modifications of the statistical quantities defined above (see SI Fig. 3 compared to Fig. 2C). However, when the exclusion radius increases to 67*nm*, deviations started to appear (SI Fig. 4). To conclude, an exclusion radius of the order of 40 *nm*, also used in [30], is consistent with the physical crowding properties of condensin and cohesin [31] to fold and unfold chromatin. Thus, using the present RCL polymer models we will now reconstruct statistical properties of chromatin in different cellular differentiation phases.

**FIG. 3.**
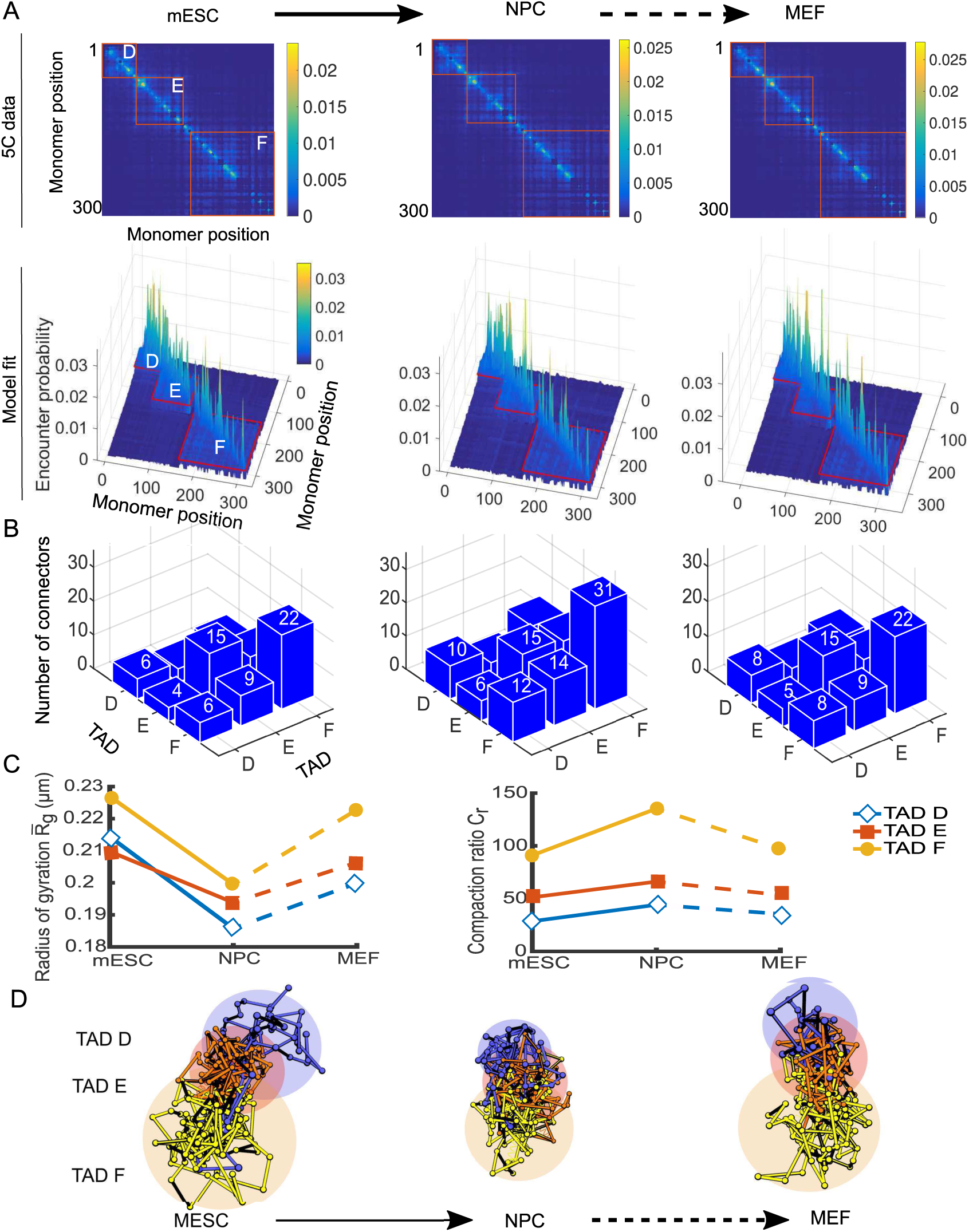
Reorganization of the X Chromosome during differentiation. **A.** 5C encounter maps and result of fitting expressions 10 and 12 to the empirical 5C encounter probability data [1] at a scale of 6kb for 3 TADs (red rectangles) at three stages of cell differentiation: mouse Embryonic stem cell (mESC, left), Neuronal precursor cells (NPC, middle) and Embryonic Fibroblasts (MEF, right). **B.** Mean number of connectors in each TAD, obtained by fitting the empirical EP as described in panel A, within and between TAD D, E and F for mESC (left), NPC (middle) and MEF (right). The number of connectors within TAD F grows from 22 for MSEC to 31 for NPC and drops back to 22 for MEF cells. The inter-TAD F and E connectivity increases from 9 for MSEC to 14 for NPC and is 9 for MEF stage. The number of connectors within TAD E remains 14 throughout the three stages, whereas the number of connectors for TAD D increases from 6 for MSEC to 10 for NPC and decreases to 8 for MEF. **C.** Mean radius of gyration (MRG, left) for the three TADs at three stages of differentiation shows synchronous compaction in the transition from mESC to NPC. The compaction ratio *C*_*r*_ (right) is computed as the cube ratio of 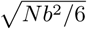 (MRG of the Rouse chain) to the MRG of each TAD with the same number of monomers *N*, shows that the compaction in TAD E is higher than that of D at NPC stage despite having a higher MRG (0.181 and 0.19*µm* for TAD D and E, respectively). **D.** Three realizations of the RCL polymer showing the compaction of TAD D (blue), E (red) and F (orange) during the transition from mESC to NPC. Shaded areas represent a ball with of gyration radius listed in panel C for each TAD in each stage.

### B. Reconstructing genome reorganization with multiple TADs from 5C data during cellular differentiation

To extract chromatin statistical properties, we constructed systematically an RCL model from 5C data of the X chromosome [1]. We focus on the chromatin organization during three stages of differentiation: undifferentiated mouse embryonic stem cells (mESC), neuronal precursor cells (NPC), and mouse embryonic fibroblasts (MEF). We first used the average of two replica of a subset of 5C data generated in [1], and then each replica separately. The two replica harbor three TADs: TAD D, E, and F, which span a genomic section of about 1.9 Mbp. We coarse-grained the 5C encounter frequency data at a scale of 6 *kb* (Fig. 3A Upper), which is twice the median length of the restriction segments of the HindII enzyme used in producing the 5C data [1, 8, 9]. At this scale, we found that long-range persistent peaks of the 5C encounter data are sufficiently smoothed out to be able to use expressions 10 and 12 for fitting the 5C EP using standard norm minimization procedure. The result is a coarse-grained encounter frequency matrix that includes pairwise encounter data of 302 equally-sized genomic segments. To determine the position of TAD boundaries, we mapped the TAD boundaries reported in bps (see [1] SI,’Analysis of 5C data, section) to genomic segments after coarse-graining. We then constructed a heterogeneous RCL polymer with *N*_*D*_ = 62, *N*_*E*_ = 88, *N*_*F*_ = 152 monomers for TAD D, E and F, respectively. To compute the minimum number of connectors within and between TADs, we fitted the EP of each monomer in the coarse-grained empirical EP matrix using formulas 10 and 12. In Fig. 3A (bottom), we present the fitted EP matrices for mESC (left) NPC (middle) and MEF (right).

We computed the number of connectors *N*_*c*_, within and between TADs, by averaging the connectivity values *ξ*_*m*_ for monomers in each TAD obtained from fitting the EP (Eq. 10 and 12) to all 302 monomers. After averaging we obtained the connectivity matrix Ξ, and used it in relations Eqs. 3–4 to recover the number of connectors within and between TADs. The mean number of connectors in the differentiation from mESC to NPC showed an increase by 145-200% (Fig. 3B) within and between TADs. The number of connectors within TAD F increases by 145% from 22 for mESC to 31 in NPC, the inter-connectivity between TAD F and E increases by 150% from 9 at mESC to 14 for NPC and the connectivity between TAD F and D doubled from 6 connectors for mESC to 12 for NPC, whereas the number of connectors within TAD E remained constant of 15. In MEF stage the number of connectors within TADs D, E and F returned to values comparable to mESC, whereas the inter-TAD connectivity between TAD F and E was 9 for MEF.

To evaluate the size of the folded TADs, we computed the MRG of the three TADs throughout differentiation stages. The MRG is the square root of the MSRG (Eq. 8) for each TAD using the calibrated connectivity matrix Ξ, obtained from fitting the experimental 5C EP (Fig. 3C, left). We found that the MRG can both increase and decrease depending on the number of connectors within TADs, but is also affected by inter-TAD connectivity, as revealed by Eq. 8. The MRG of all TADs decreased in average from 0.21*µm* at mESC stage to 0.19*µm* for NPC and was 0.2*µm* for MEF cells. From mESC to NPC, the MRG of TAD E exceeded that of TAD D (Fig. 3C, red squares) despite the higher numbers of added connectors in TAD D and its smaller size *N*_*D*_ = 66 compared to *N_E_* = 88. This result shows how the inter-TAD connectivity contributes to determine the MRG and the volume of TADs. In addition, using the calibrated RCL model at 3kb, we were able to reproduce the distributions of three-dimensional distances between seven DNA FISH probes reported in [8] (SI Fig. 2).

Finally, we recall that a ball of radius of gyration is insufficient to characterize the degree of compaction inside a TAD, because it does not give the density of bps per *nm*^3^. To obtain a better characterization of chromatin compaction, we use the compaction ratio for TAD_*i*_, defined by the ratio of volumes:

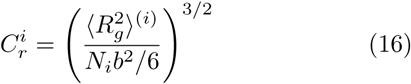

where 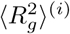 is given by formula 8 and the denominator is the MSRG for a linear Rouse chain of size *N*_*i*_[13]. We find that TAD F (*N*_*F*_ = 154 monomers) has the highest compaction ratio among all the TADs among the three stages of differentiation (Fig. 3C, circles): indeed for TAD F, 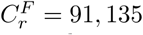, 135, and 97 fold more compact than the linear Rouse chain with *N* = 150 monomers, associated with mESC, NPC, and MEF stages, respectively. For TAD E, (*N*_*E*_ = 88 monomers) the compaction ratio is 51, 66, and 45, thus it is more compact than the linear Rouse chain with *N* = 88 monomers (Fig. 3C right, red squares), despite retaining 15 intra-TAD connectors in all stages of differentiation (panel B). This effect is due to an increased inter-TAD connectivities between TAD E and F at NPC stage to 15. Finally, TAD D (*N* = 62 monomers), characterized by 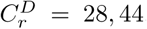, and 35 (blue diamonds) is more compact than a Rouse chain of *N* = 62 monomers, for mESC, NPC and MEF stages, respectively.

To examine the consistency of our approach and the ability of RCL model to represent chromatin, we fitted independently the EPs *P*^(1)^, *P*^(2)^ of the 5C data of replica 1 and 2 at 10 kb resolution (SI Fig. 5A-C) using Eqs. 10–12 (Methods). We found that the number of added connectors in replica 1 and 2 differs by at most five connectors for TAD F. This difference between replica may arise from intrinsic fluctuations in the statistics of encounter frequencies. We further compared the EP *P*^(1)^ with the empirical EP *E*^(2)^ of replica 2 (SI Fig. 5D, left); We found that 〈||*P*^(1)^(*m*) − *E*^(2)^(*m*)_*m*_||〉_*m*_, averaged over monomers *m* (SI, Eq. 65), equals 0.17. Note, that the main contribution of this difference arises from monomers forming long-range loops (SI Fig.5D) consistent with off-diagonal peaks of the 5C data. Similarly, we found 〈||*P*^(2)^ − *E*^(1)^||〉_*m*_ = 0.17 (SI Fig.5D, right). In addition, the mean radii of gyration for all three TADs in both replicas were comparable for all three stages of differentiation (SI Fig. 6A-B, left).

To evaluate the consequences of boundaries between TADs on the number of added connectors necessary to reconstruct the heterogeneous RCL polymer (SI Fig. 7), we subdivided each TAD D-F into two equal parts, and repeated the fitting of the heterogeneous RCL model to the EP of the six resulting sub-TADs. We tested the scenario where the number and boundaries of TADs differ from the one in [1], which can result from various TAD-calling algorithms [32, 33]. We extracted the intra and inter-TAD connectivity fractions Ξ and computed the number of connectors by fitting the empirical EP (SI Fig. 6A-B). We computed the difference in the average number of connectors between the three TADs case (SI Fig. 6C) and the six sub-TADs case. We found a maximal difference of six connectors for intra-TAD connectivity of TAD E in the MEF stage (SI Fig. 7D). For inter-TAD connectivity, the average difference is two connectors. In addition, we find that the compaction of TADs throughout differentiation is preserved for the six TAD case (SI Fig. 8A), and further find a qualitative agreement between the compaction ratios of the three and six TADs case (see comparison, SI Fig. 8B).

Finally, to determine the robustness of the predictions of the heterogeneous RCL polymer, we compared the reconstructed 5C statistics (Fig. 3) to the statistics reconstructed from HiC data [34] of the X chromosome, harboring TAD D, E, and F, binned at 10kb, with *b* = 1.814*µm* computed from SI Eq. 67, and for three successive stages of differentiation: mESC, NPC, and cortical neurons (Fig. 4). Note that the polymer model reconstructed from HiC and 5C data, are not necessary identical, although their share some similar statistics, because for each one, the data are generated at a different resolution. However, we found a good agreement between the intra-TAD connectivity of TADs D and E of the 5C and HiC data for mESC and NPc stages (Fig. 4A, C). In general, the inter-TAD connectivity in the HiC data was lower (average of 1.5 connectors) than that of the 5C (average of 4), which resulted in an increased MRG for all TADs (Fig. 4B, D, left and a decreased compaction ratios 4B, D, right). A direct comparison between the reconstructed statistics of the 5C MEF and HiC CN was not possible. To conclude, inter-TAD connectivity plays a key role in the compaction of TADs and therefore recovering their exact number is a key step for precisely recovering genome reorganization from 5C data.

**FIG. 4.**
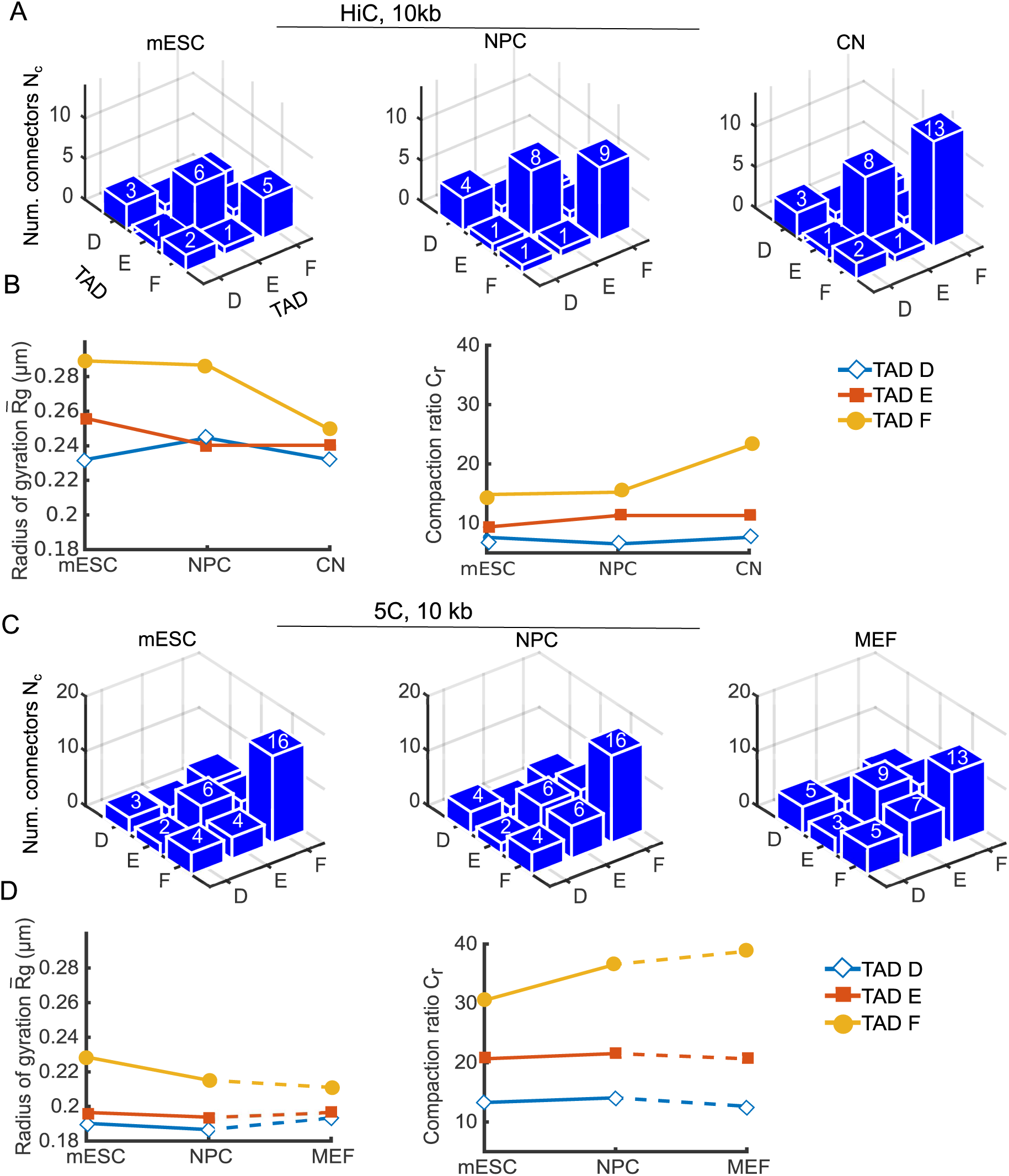
Comparing TADs reconstruction during cell differentiation between HiC and 5C. **A.** Average number of connectors within and between TADs D, E, and F of HiC data [34] of the X chromosome binned at 10 kb, obtained by fitting the empirical EP with Eq. 10, 12, where TAD boundaries were obtained in [1] for mouse embryonic stem cells (mESC, left), neuronal progenitor cells (NPC, middle), and cortical neurons (CN, right). The average number of connectors within and between TADs are presented in each blue box. **B.** Mean radius of gyration (left) for TAD D, E, and F, throughout three successive stages of differentiation of the HiC data, with *b* = 0.18*µm* obtained from SI Eq. 67, and the compaction ratio (right, Eq. 16). **C.** Average number of connectors within and between TADs D, E, and F of the 5C data [1] of the X chromosome binned at 10 kb, obtained by fitting the empirical EP with Eq. 10, 12 for mESC (left), NPC (middle), and MEF (right). **D.** Mean radius of gyration (left) for TAD D, E, and F of the 5C data, and the compaction ratio (right, Eq. 16).

### C. Distribution of anomalous exponents for single monomer trajectories

Multiple interacting TADs in a cross-linked chromatin environment, mediated by cohesin molecules can affect the dynamics of single loci trajectories. Indeed, analysis of single particle trajectories (SPTs) [35–39] of a tagged locus revealed a deviation from classical diffusion as measured by the anomalous exponent. We recall briefly that the MSD (Eq. 14) is computed from the positions *r*_*i*_(*t*) of all monomers *i* = 1, …, *N*_*T*_. In that case, the MSD, which is an average over realization, behaves for small time t, as a power law

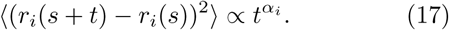

It is still unclear how the value of the anomalous exponent *α*_*i*_ relates to the local chromatin environment, although it reflects some of its statistical properties, such as the local cross-link interaction between loci [20, 38]. Thus we decided to explore here how the distribution of cross-links extracted from EP of the HiC data could influence the anomalous exponents. For that purpose, we simulate a heterogeneous RCL model, where the number of crosslink was previously calibrated to the data. The number and position of the connectors remain fixed throughout all the simulations (for tens of seconds).

We started with a heterogeneous RCL model with three TADs, reflecting the inter and intra-TAD connectivity as shown in Fig. 3. We generated a hundred chromatin realizations 𝒢_1_, …, 𝒢_100_. In each realization 𝒢_*k*_, the position of added connectors is not changing. We then simulated in time each configuration a hundred times until relaxation time (SI Eq. 23). After the relaxation time is reached, defined as *t* = 0, we followed the position of each monomer and computed the MSD up to time *t* = 25*s*. To compute the anomalous exponent *α*_*i*_, we fitted the MSD curves using a power law Eq. 17 to estimate the anomalous exponents *α*_*i*_, *i* = 1, …, 302 along the polymer chain. We repeated the procedure for each stage of cell differentiation: mESC, NPC, and MEF.

In Fig. 5, we plotted the anomalous *α*_*i*_ for each monomer of the three stages mESC (left), NPC (middle), and MEF (right), and for TAD D (dark blue), TAD E (cyan) and TAD F (brown). We find a wide distribution of *α*_*i*_ with values in the range *α*_*i*_ ∈ [0.25, 0.65] for all TADs in the three cell types. The average anomalous exponent in TAD D is *α*_*D*_ = 0.46, in mESC stage, reduced to *α*_*D*_ = 0.41 in NPC, due to the increases intra-TAD connectivity, and increased to *α*_*D*_ = 0.435 in NPC stage. The average anomalous exponent in TAD E, *α*_*E*_ =0.425, 0.41, 0.426 at mESC, NPC, and MEF stages, respectively. The average anomalous exponent of TAD F was *α*_*F*_ =0.443, 0.405, 0.44 at mESC, NPC and MEF stages, respectively. The anomalous exponent *α* decreases with adding connectors, observed throughout differentiation in all TADs, which is in agreement with the compaction and decompaction of TADs (Fig. 3C and SI Figs. 6–7). Furthermore, we obtain an average anomalous exponent of 0.4, previously reported experimentally in [35]. To complement the anomalous exponent, we estimated the space explored by monomers by computing the length of constraint *L_c_* [38] (computed empirically along a trajectory of *N_p_* points for monomer R as 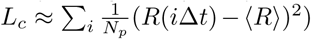 for three monomers in each TAD D, E, F: *r*_20_, *r*_70_,*r*_120_. For a single connector realization, we obtain *L*_*c*_ ≈ 0.3, 0.25,0.26*μm*, respectively, which is about twice the simulated MRG of TAD D, E, F: 0:18; 0:13:0:17*μm*, respectively. Thus we conclude that random distributions of fixed connectors can reproduce the large variability of anomalous exponents reported in experimental systems using single locus trajectories, especially for bacteria and yeast genome [35, 38] in various conditions.

**FIG. 5.**
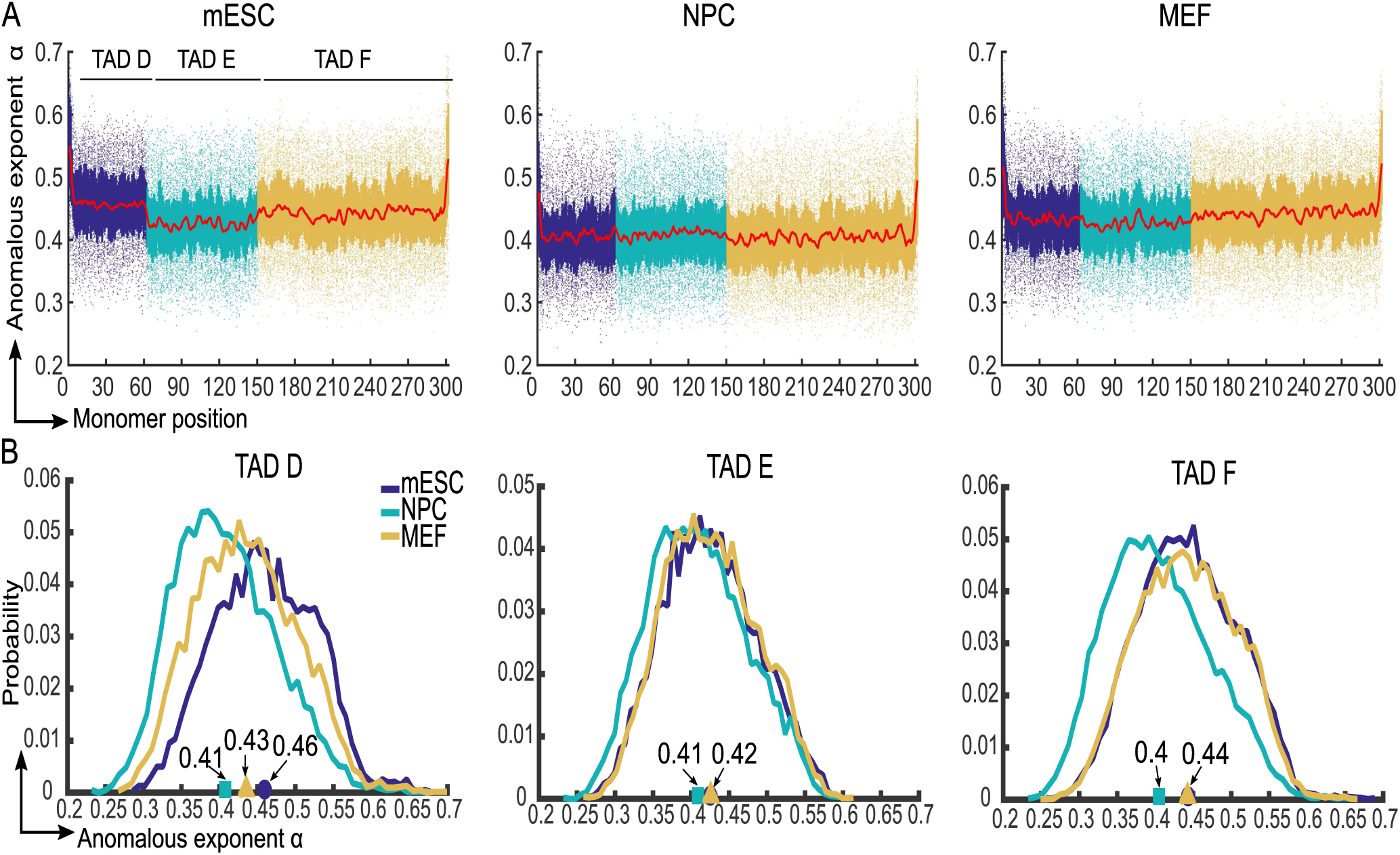
Anomalous exponents in three stages of differentiation. **A.** Anomalous exponents computed for 302 monomers of the RCL polymer reconstructed in Fig. 3, corresponding to 5C data of three TADs, TAD D (dark blue), TAD E (cyan), and TAD F (brown) of the X chromosome [1]. A hundred realizations are simulated for each configuration using Eq. 7. For each realization, we choose the positions of added connectors uniformly distributed within and between TADs and repeated simulations 100 times, after relaxation time has been reach(SI Eq. 23). We then run simulation for 25*s*. The anomalous exponents *α*_*i*_, *i* = 1, …, 302 are obtained by fitting the MSD curve of each monomer using model 17. **B.** Distribution of the anomalous exponent in TAD D (left) TAD E (middle) and TAD F (right) for three cell stages: mESC (dark blue), NPC (cyan), and MEF (brown). The average anomalous exponents in TAD D, E, and F, are *α*_*D*_ =0.46, 0.41, 0.435, *α*_*E*_ =0.425, 0.41, 0.426, and *α*_*F*_ =0.443, 0.405, 0.44 for mESC (circle), NPC (square), and MEF (triangle) stages, respectively.

## DISCUSSION

We presented here a general framework based on the RCL model [9, 11] to extract statistical and physical properties of multiple interacting TADs. The present polymer model differs from others by several aspects: Our construction of a polymer model from HiC is parsimonious.

It uses a minimal number of added connectors at a given scale to match the experimental steady-state of HiC/5C data, in contrast to the model [8], which is based on a full monomer-monomer interaction, described by pairs of potential wells. In addition, at the scale of few *µm* occupied by TAD D, E and F of chromosome X, we neglected crowding effects from neighboring chromosomes. Furthermore, we do not use here several types of diffusing binders that need to find a binding site in order to generate a stable link, as introduced in [40]. In addition, contrary to the Random Loop Model (RLM) [16], we do not consider here transient binding, because the positions of random connectors within TADs does not matter (as long as they are uniformly randomly placed). By placing connectors randomly, we capture the heterogeneity in chromatin organization across cell population. Here we fix connectors, which are stable in the time scale of minutes to hours. The present polymer construction is motivated by the evidence of many stable loci-loci interactions, which are common to the majority of chromosomes in 5C (e.g. peaks of the 5C data) [1, 33]. These stable interactions are also found at TAD boundaries, which are conserved in both human and mouse. There are several conflicting studies ([5, 33, 41]) about the binding time of connectors (CTCF-cohesin, etc…), which suggest that cross-links can remain stable for minutes to hours and even during the entire phase cycle. Here, we study the chromatin dynamics within this time range where cross-links are stable [33, 41]. One final difference between the present RCL model and the RLM model, is the possibility to account for several interacting TADs and our new expressions for statistical quantities such as the radius of gyration, encounter probability or MSD.

We applied our framework to reconstruct chromatin organization across cell differentiation from conformation capture (3C, 5C, and Hi-C) data, where we accounted for both intra and inter-TAD connectivities. The RCL polymer model allows estimating average number of cross-links within and between each TAD, length-scales such as the mean square radius of gyration that characterizes the size of folded TADs, and the mean square displacement of monomers in multiple interacting TADs. The present method allowed us to estimate the volume occupied by TADs. These quantities cannot be derived directly from the empirical conformation capture data and are usually extracted from SPT experiments [42, 43]. Finally, the present approach can also be used to study the phase-liquid transition at the chromatin level, depending on the number of connectors. We computed here the Radius of Gyration that clarifies how chromatin compaction depends on the number of added connectors. Similar questions were recently discussed in a general perspective [44] and we presented here some answers.

We have applied the present approach to reconstruct multiple TAD reorganization during three stages of cell differentiation: mouse embryonic stem cell (mESC), neuronal precursor cells (NPC), and mouse embryonic fibroblasts (MEF). We fitted expressions 10 and 12 to the empirical 5C encounter matrices of three differentiation stages (Fig. 3A) and showed that the RCL model produced contact enriched TADs with a variability in monomer connectivity within each TAD (Fig. 3A, super and sub-diagonals of EP matrices). We use the average connectivity obtained to compute the average number of connectors within and between TADs (Fig. 3B). At a scale of 6 kb, our results show that the X-chromosome acquires connectors within and between TADs D, E, and F in the transition between mESC to NPC, and the number of connectors is comparable to those of mESC at MEF stage. Increased connectivity for NPC cells is correlated with an increase of LaminB1 (see [1], Figure 3). Similarly, we reported (Fig. 3A-B) an increase in the number of connectors within TADs D, E, and F in NPC stage in comparison to the mESC stage, which is associated with TAD compaction. Indeed, the MRG curves (Fig. 3C, left) decrease from mESC to NPC stage, indicating a higher chromatin compaction for all TADs. The compaction ratio (compared to Rouse polymer) showed a high compaction at NPC stage (Fig. 3C, right) for all TADs, which can be associated with a heterochromatin state, suppressed gene expression and lamina associating domains [45]. Inter-TAD connectivity remained quite stable as the number of connectors for TAD E did not change, but the MRG decreased and the compaction ratio increased at NPC stage. This result shows that an accurate description of chromatin from 5C data by polymer models has to account for inter-TAD connectivity. Indeed, despite having similar intra-TAD connectivity, the reconstruction of 5C and HiC differs in inter-TAD connectivity, which is reflected in a change in the MRG (Fig. 4). In addition, using Eq. 15, we were able to reproduce the distribution of three-dimensional distances between seven genomic loci, measured by DNA FISH probes (SI Fig. 2). Overall, the RCL polymer captures the correlated reorganization of TAD D, E, and F during differentiation. TAD reorganization is also compatible with transcription co-regulation in these TADs [1]. Moreover, we found here multiple connected regions with weak inter-TAD connectivity, as suggested by experimental conformation capture data [1, 4, 5, 12] that can affect all loci dynamics inside a TAD (Fig. 5).

To describe 5C encounter probability of HiC data, we used theory (Eqs. 10 and 12) and simulations so we could capture both TADs and higher order structures (meta-TADs [12]) resulting from weak inter-TAD connectivity (Fig. 2A and B). This representation allowed us to clarify how the dynamics of monomers are affected by the local connectivities within and between TADs [46]. Using the MSD curves (Fig. 2C), we found that local connectivities (number and positions) are responsible to shape the value of the anomalous exponents, leading to a large spectrum with a mean of 0.4, (Fig. 5). This result explains the large variability in MSD and anomalous exponent behavior reported experimentally in SPT experiments [9, 18, 35, 37, 42, 47–49]. Indeed, the local chromatin organization can vary in cell population, as cohesin could bind randomly at various places. Furthermore, we have shown here that the anomalous exponents are modulated by the number and the positions of linkers. The value of anomalous exponent *α* does not depend in general on the diffusion coefficient *D* or *b*, as known in various other polymer models, such as for Rouse or *β*-polymers [20, 50]. We have shown here (Fig.5) that the anomalous exponent crucially depends on the number and the distribution of connectors (see also Fig. 3D). In the mean-field approximation, the mean exponent is 0.5, which does not depend on the polymer scale. Finding the exact relation between the number of connectors for a specific connector configuration (not in the mean-field case) remains challenging and relevant to reconstruct the local connectors environment from measured anomalous exponent. To conclude, we propose that measuring the anomalous exponents of loci, positioned at different locations inside a TAD, could reveal the amount of connectors and thus chromatin condensation beyond the exact position of these loci. In summary, the present method allows to reconstruct a polymer model from HiC, to generate numerical simulations and to estimate the MSD and the anomalous exponents, relating HiC with SPT statistics.

We emphasize here that the present approach can be used to describe chromatin with volume exclusion lower than 40*nm* (SI Fig. 3–4. Furthermore, we obtained here estimations for the number of connectors within and between TADs, which was consistent across two replicas of the 5C data (SI Fig. 7–8). Comparing the predicted number of connectors between 5C and HiC data, we found a good agreement in the intra-TAD connectivity. However, the inter-TAD connectivity was reduced in the HiC vs. 5C (Fig. 4). We attributed these discrepancies to the elimination of inter-TAD interactions, which are smoothed out at 10 kb resolution for the HiC data. Connectors are likely to represent binding molecules such as cohesin and the present results suggest that few of them are actually needed to condense chromatin and their exact positions inside a TAD does not necessarily matter. The interpretation of the number of connectors *Nc* is not straightforward. This number characterize the amount of connectors at a given scale, which suggests that in the limit of 1bp resolution *Nc*, it would be equal to the endogenous number of linkers, but as the resolution decreases, the number of connectors that are reported by the HiC data should also decrease, because two connectors in the same bin are not counted (Fig. 6A). It is always possible to coarse-grain a polymer model, but reconstructing a refined polymer from a coarse grained kb-scale is an ill-posed problem, because the refined encounter probability at high kb-resolution cannot be inferred from a low resolution (yellow and red arrows in Fig. 6B).

**FIG. 6.**
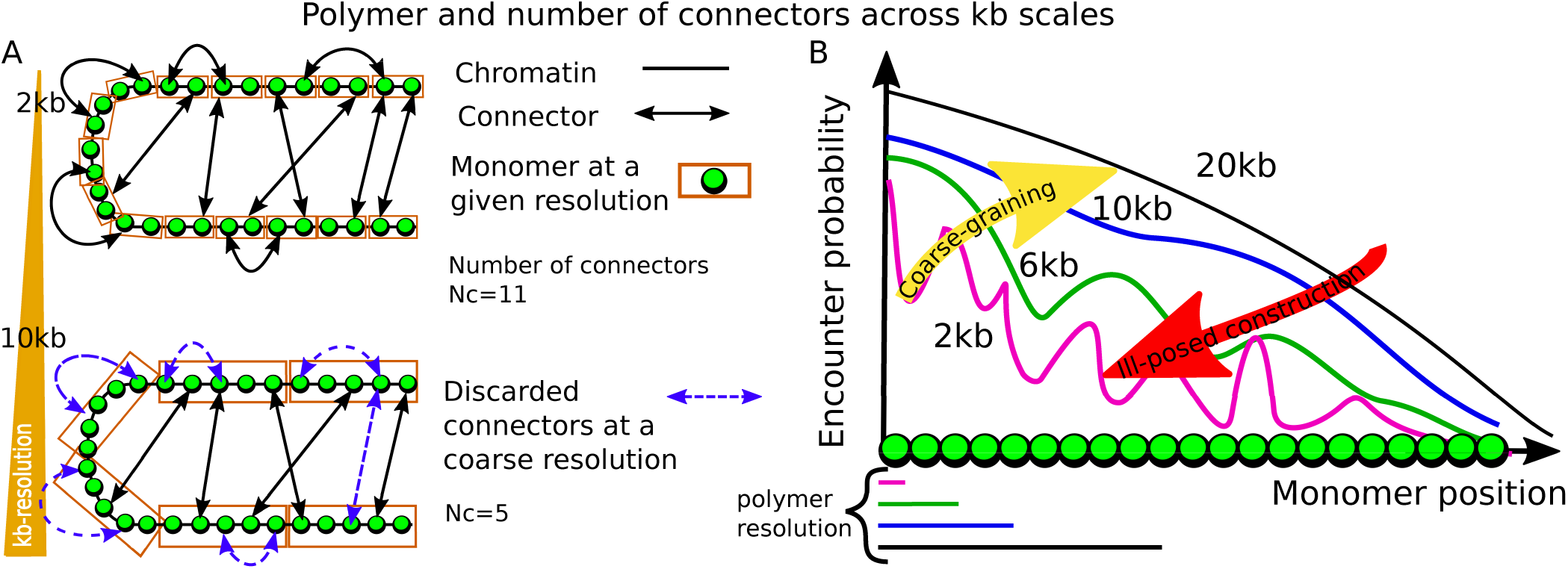
RIllustration of polymer reconstruction of chromatin at different kb-scales. **A.** Two examples of polymer reconstruction at 2 (Upper) and 10 kb (bottom) resolution, where the number of connectors *Nc* vary with the scale *Nc* = 11 (2kb) and *Nc* = 5 (10kb). For coarse scale, connectors within the same bin (orange boxes) are discarded. **B.** Effect of coarsegraining on the encounter probability. It is possible to coarse-grain (yellow arrow), while the construction of a refined polymer model from a coarse-grained resolution (red arrow) is an ill-posed problem.

To conclude, the present framework is a new tool to reconstruct systematically chromatin structural reorganization from HiC matrices and can be used to interpret chromatin capture data. In particular, using any experimental data of ligation proximity experiments (Hi-C, 5C 4C), we obtained here statistical properties beyond conformation capture data. The present analysis could also be applied to reveal the statistics of complex matter at the transition point between liquid and gel [44, 51] based on heterogeneous random architecture at the level of chromatin sub-organization.

## ACKNOWLEDGMENTS

VP and DN acknowledge funding from the Fondation pour la Recherche Médicale (AJE20140630069) and the Agence Nationale de la Recherche (ANR-14-ACHN-000901 and ANR-16-TERC-0027-01). DH’s research is supported by FMR team DEQ20160334882.

## SUPPLEMENTARY INFORMATION

### I. CONSTRUCTING A RCL POLYMER FOR A SINGLE TOPOLOGICAL ASSOCIATING DOMAIN (TAD)

The RCL polymer for a single TAD [7] in dimension *d* = 3 consists of *N* monomers at positions ***R*** = [*r*_1_, *r*_2_, …, *r*_*N*_]^*T*^, connected sequentially by harmonic springs [8]. Then *N*_*c*_ spring connectors are added randomly between non-nearest neighboring (non-NN) monomer pairs (Fig. 1A, main text) leading to a realization *G*. The energy 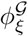 of the RCL polymer is the sum of the spring potential of the linear backbone plus that of added random connectors (see also [1, 2]):

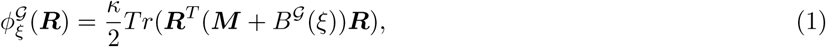

where *κ* is the spring constant, and *Tr* is the trace operator. The matrix ***M*** is the Rouse matrix for the linear backbone [8], defining connectivity between NN monomers

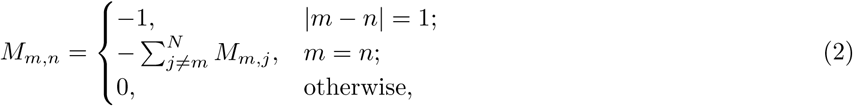

and *B*^𝒢^(ξ) is a square symmetric connectivity matrix between *N*_*c*_ randomly chosen non-NN monomer pairs,

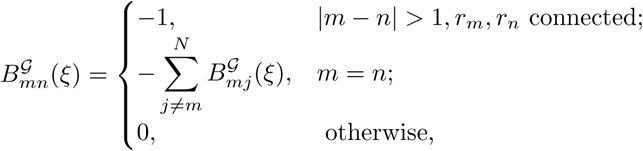

and the connectivity fraction

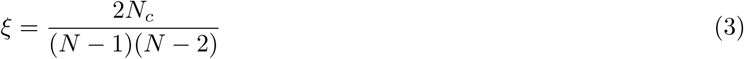

is the ratio of the number of connectors *N*_*c*_ to the maximal possible non-NN monomer pairs to connect.

The dynamics of monomers (vector ***R***) is given by the sum the field of forces obtained from the energy 1 plus Brownian motion:

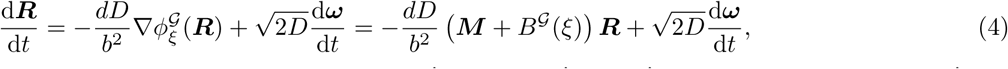

where *D* is the diffusion constant, and ***ω*** are standard *d*-dimensional Brownian motions with mean zero and variance one. To study the statistical properties of each monomer in Eq. 4, we previously used a mean-field approach [7], where we replaced the matrix *B*^*𝒢*^ (*ξ*) by its average 〈*B*^*𝒢*^ (*ξ*)〉 (average over all realizations *G* for a given fraction of connectors *ξ* [1, 7]). We computed the steady-state variance, the encounter probability (EP) between monomers *m, n* of a single TAD [7], which was used to fit 5C data of a single TAD to recover the number *N*_*c*_ of connectors.

### II. CONSTRUCTING A HETEROGENEOUS RCL POLYMER FOR MULTIPLE INTERACTING TADS

In this section we generalize the construction of the RCL polymer of a single TAD (Section I) to heterogeneous RCL polymer, consisting of multiple interacting TADs. We derive statistical properties of the heterogeneous RCL model, such as intra and inter-TAD encounter probabilities, mean radius of gyration of each TAD, and the distribution of the spatial distances between monomers.

#### A. Construction of the average connectivity matrix 〈*B*^𝒢^(Ξ)〉

**FIG.S 1.**
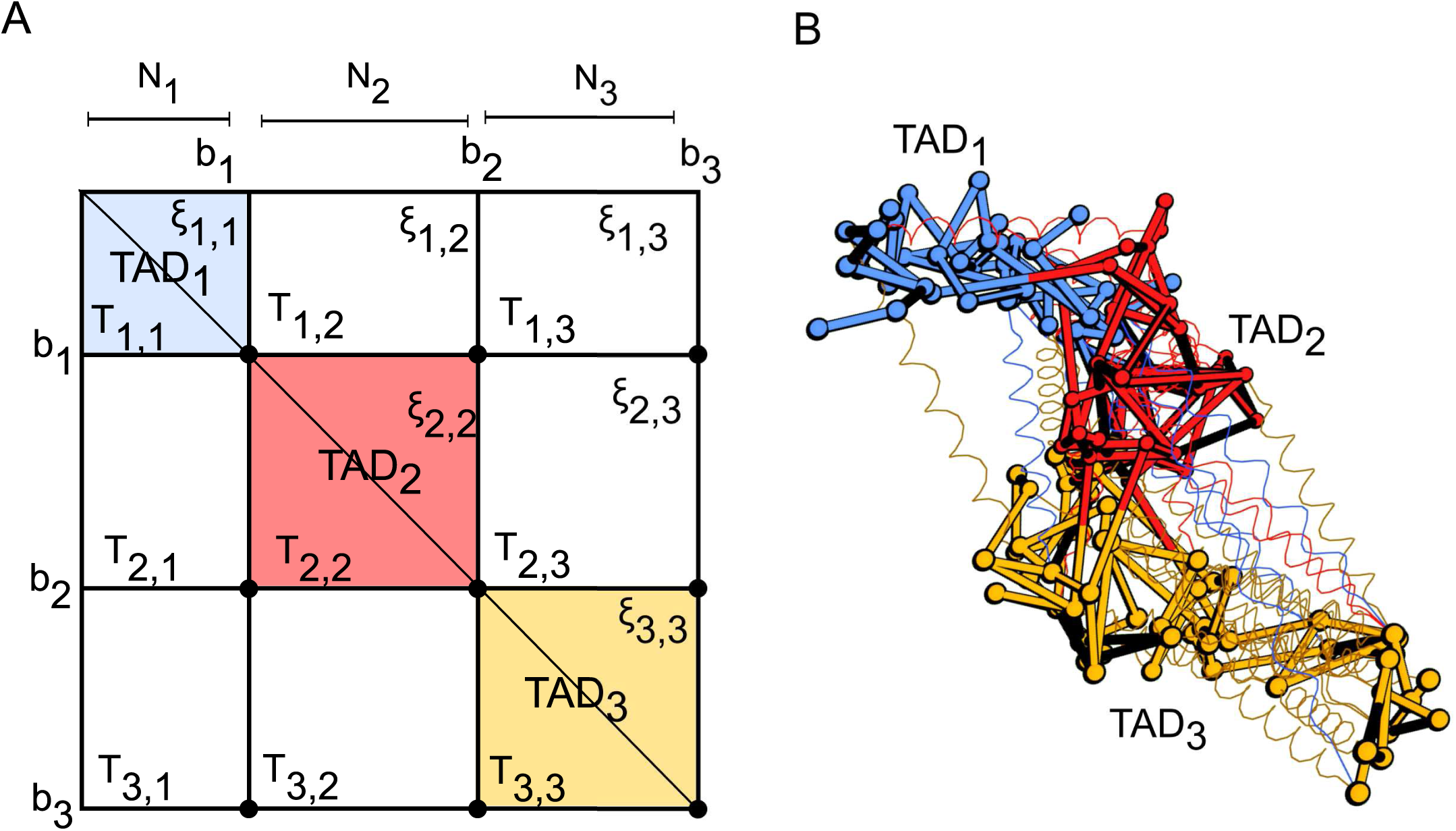
Constructing the heterogeneous RCL polymer. **A.** Schematic description of 3 TADs: TAD_1_ (blue) TAD_2_ (red) and TAD_3_ (orange) composed of *N*_1_, *N*_2_ and *N*_3_ monomers connected linearly, containing additional random connectors within and between TADs, determined by the connectivity fraction *ξ_*ij*_* (Eq. 4, main text). The indices *b*_*i*_ represent the bottom right end of each block associated to the grid points (*b*_*i*_, *b*_*j*_), which define blocks [*T*_*ij*_]. **B.** Sample configuration of a heterogeneous RCL polymer with three TADs corresponding to the construction in panel A. Monomers (spheres) are connected linearly by harmonic springs to form the backbone, and additional connectors (springs) are added randomly between non-nearest neighboring monomers within and between TADs.

To construct the average connectivity matrix

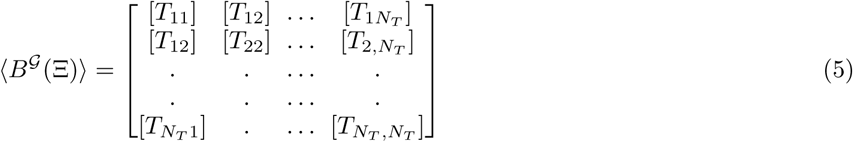

(see Fig. S1A), we start with matrix *B*^*𝒢*^ (Ξ) (see Eq. 5, main text) and define boundaries of blocks [*T*_*ij*_], 1 ≤ *i, j* ≤ *N*_*T*_ : the upper left and lower right indices of block [*T*_*ij*_] are (*b*_*i*−1_, *b*_*j*−1_) and (*b*_*i*_, *b*_*j*_), respectively, where

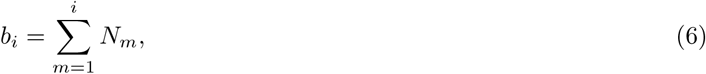

and *b*_0_ = 1.

We shall now describe the procedure of averaging. For that purpose, we chose *N*_*c*_(*i, j*) independent pairs of monomers between blocks [*T*_*ij*_], (*i* ≠ *j*) with a uniform probability, where *N*_*c*_(*i, j*) is the number of connected pairs between TAD *i* and *j*. By symmetry, we can restrict the choice of monomer pairs to the upper triangle of 〈*B*^*𝒢*^ *(Ξ)〉*. We now compute the probability 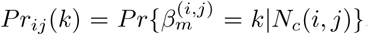, of monomer *m* in TAD *i* (*m* ∈ (*b*_*i*‒1_, *b*_*i*_)), to be connected to *k* non-NN monomers in TAD *j*, given the total number of connectors *N*_*c*_(*i, j*). For *i* = *j*, inside TAD *i*, this probability is computed by choosing *k* indices in row *m* and the remaining *N*_*c*_(*i, i*) − *k* connectors in any row or column *n* ≠ *m*, leading to the hyper-geometric probability

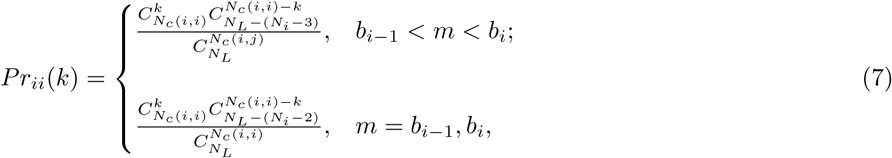

where 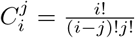 is the binomial coefficient.

For off-diagonal blocks [*T*_*ij*_], with *j > i*, the probability of monomer *m* to be connected to *k* monomers is

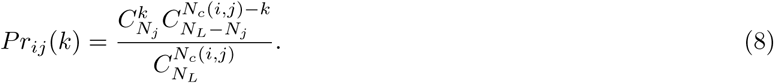

The mean number of connectors 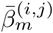, for monomer *b*_*i−*1_ < *m* < *b*_*i*_ in each block [*T*_*ij*_] is computed from formulas 7-8

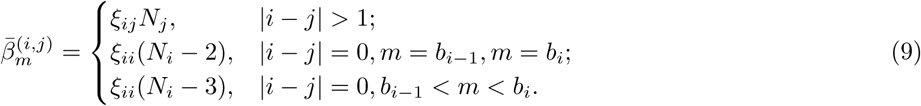

We now compute the average number of connectors for monomer *m*. This computation is obtained by the sum of average intra and inter-TAD connectors (Eq. 9) over all blocks [*T*_*ij*_], *j* = 1, .., *N*_*T*_

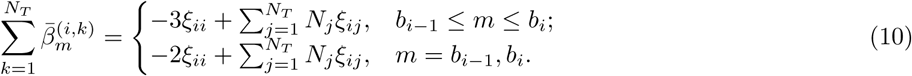

We can therefore represent each block [*T*_*ij*_] in a matrix form by

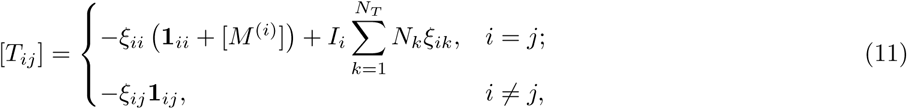

where **1**_*ij*_ is a *N*_*i*_ × *N*_*j*_ matrix of ones, [*M*^(*i*)^] is a Rouse matrix (Eq. 2) for a polymer of *N*_*i*_ monomers, and *I*_*i*_ is an *N*_*i*_ × *N*_*i*_ identity matrix. This ends the computation of the average connectivity matrix 〈*B*^*𝒢*^ (Ξ)〉 defined in Eq. 5 by blocks 11.

The energy of the mean-field heterogeneous RCL polymer is given by Eq. 1, with the connectivity fraction matrix Ξ (Eq. 2, main text), and average random connectivity matrix 〈*B*^*𝒢*^ (Ξ)〉 (Eq. 5). The mean-field equation describing the dynamics of monomers ***R*** is

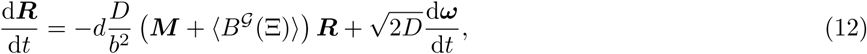

where *d* is the dimension, *b* is the standard-deviation of connected monomer for *Nc* = 0 (Rouse chain), *M* = *diag* [*M*^(1)^], …, [*M* ^(*N*_*T*_)^], and *ω* are Brownian motions with mean zero and standard-deviation one.

#### B. Relaxation time for the average RCL polymer

We derive now the statistical properties of the monomer dynamics based on Eq. 12. We first diagonalize the average connectivity matrix 〈(*B*^*𝒢*^ (Ξ) 〉 (drift term) using the normal coordinates transformation

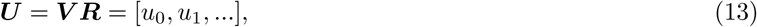

where

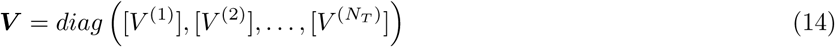

is a diagonal block matrix. Each block [*V* ^(*i*)^] is a *N*_*i*_ × *N*_*i*_ matrix of eigenvectors associated with the Rouse polymer containing *N*_*i*_ monomers, and is given by

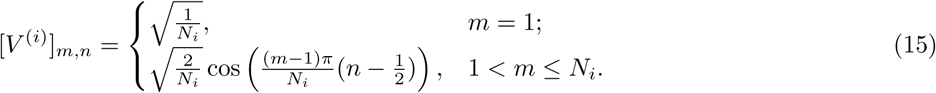

To obtain the normal form, we multiply 12 from the left by 14 and obtain

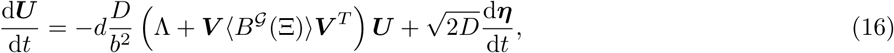

where ***η*** = ***V ω*** are standard Brownian motions with mean zero and standard-deviation one. The *N*_*T*_ } *N*_*T*_ block diagonal matrix

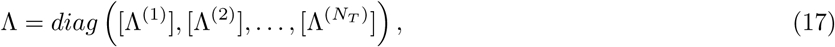

consists of eigenvalues of Rouse chains, containing *N*_1_, *N*_2_, …, *N*_*T*_ monomers, and is defined by

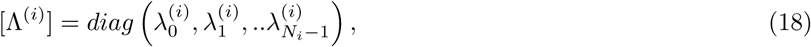

where the Rouse eigenvalues are

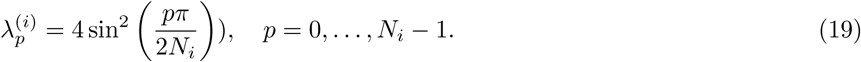

To compute ***V*** 〈*B*(Ξ)〉 ***V***^*T*^ in Eq. 16, we use relation 11, and note that [*T*_*ii*_] commutes with [*M* ^(*i*)^] (Eq. 2), and therefore we can carry the computation for each block separately to obtain

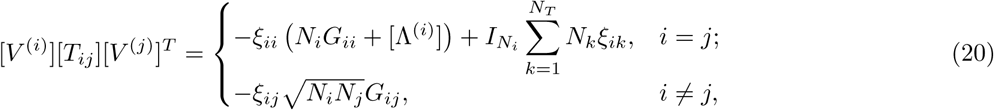

where *G*_*ij*_ (*m, n*) is a *N*_*i*_ × *N*_*j*_ matrix made of zeros except for *G*_*ij*_(1, 1) = 1. Thus, the normal coordinate 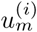 (1 ≤ *m* ≤ *N*_*i*_ − 1) in chain *i* satisfies

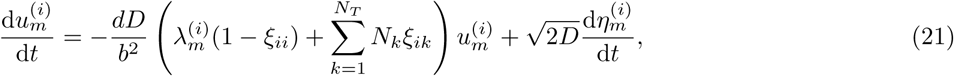

whereas the centers of masses 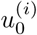 are driven by

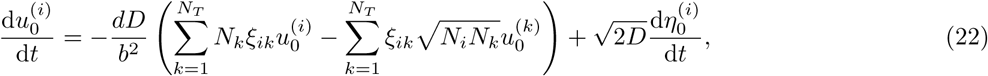

where 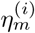 are *d*-dimensional Brownian motions with mean zero and standard-deviation one. To conclude, the relaxation time for a polymer containing *N*_*T*_ TADs is computed from of Eq. 21 as the slowest relaxation time

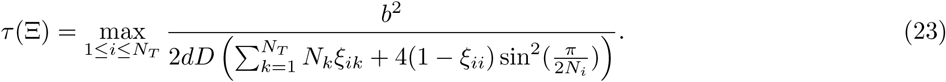

Using the values *b* = 0.2*µ, D* = 0.008*µ*^2^/*s*, and *Nc* for polymers as described in Fig. 3 and 5, and Eq. 23, we computed numerically the relaxation times for TAD D, E and F and found that *τ*_*D*_ ≈ 28*s*; *τ*_*E*_ ≈ 12*s*; *τ*_*F*_ ≈ 16*s*. All statistical quantities were computed here from simulations after we waited 10,000 time steps, which is much more than these relaxation times (hundreds of seconds), ensuring that the polymer model has reached a steady-state. Note that the internal monomer position fluctuations due to thermal noise contributes to the dynamics of each monomer, captured by the MSD, which is a second order statistical moment.

#### C. Mean square radius of gyration for each TAD

The mean square radius of gyration (MSRG) characterizes the average size of a folded TAD. We derive now an expression for the probability distribution of the square radius of gyration of each TAD across realizations *G* and then compute the first two moments. We start with the formula for the distribution of the square radius of gyration [11]

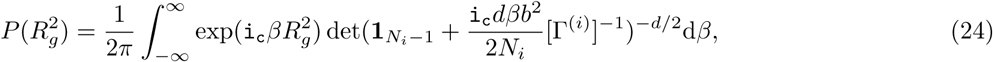

where i_c_ is the complex unit, [Γ^(*i*)^] is a (*N*_*i*_ − 1) × (*N*_*i*_ − 1) diagonal matrix of eigenvalues for the block connectivity matrix [*M*^(*i*)^] + [*T*_*ii*_] (Eqs. 2 and 11), excluding the first zero eigenvalue, and det is the determinant operator. We evaluate now the determinant in 24 by a series expansion

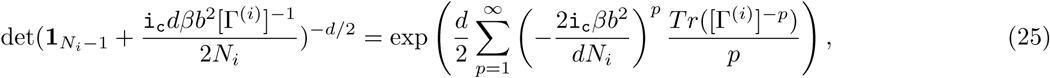

where *Tr* is the trace operator. We truncate the series in 25 at order *p* = 2 and substitute the result in 24 to obtain

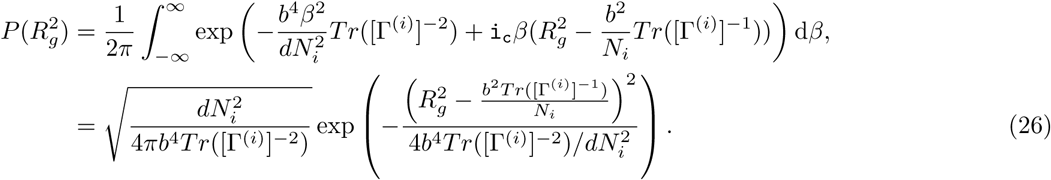

Thus, the distribution of the square radius of gyration of each TAD_*i*_ is Gaussian with mean and variance given respectively by

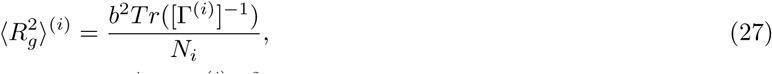

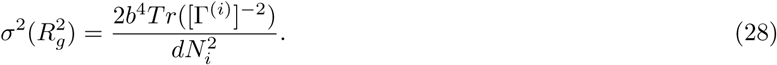

Under the assumption of dominant intra-TAD connectivity,

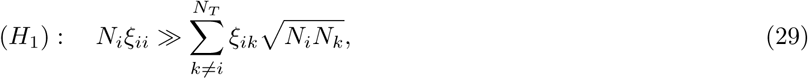

which is well satisfied for TADs maps [3–5], we compute Eq. 27 by first approximating

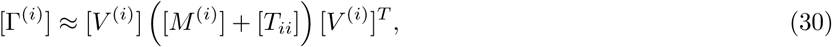

which is valid for weak inter-TAD connectivity (assumption Eq. 29), where the center of masses of TADs are decoupled in Eq. 22. We then use the approximation 30 for *N_i_ »* 1 to evaluate the trace in 26

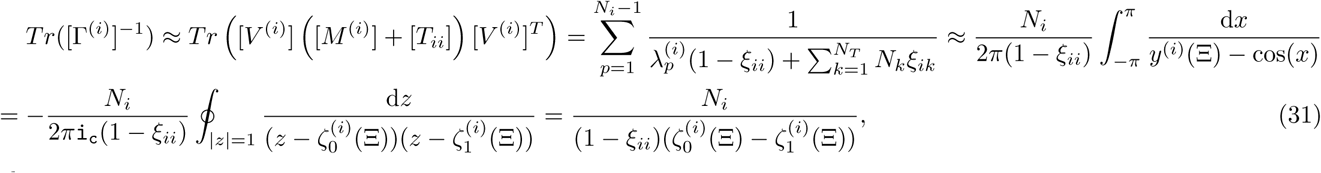

where

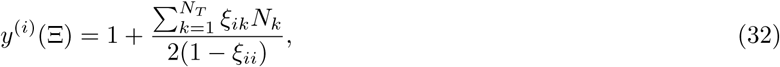

and 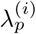 (Eq. 19) are the eigenvalues of the Rouse block matrix (Eq. 2) and

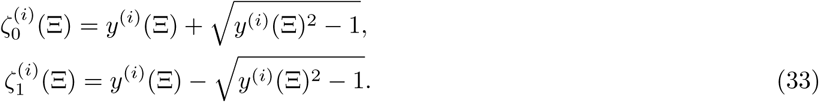

Substituting Eq. 32 and 33 into 31 and then into 27, we obtain the expression for the MSRG of TAD *i*

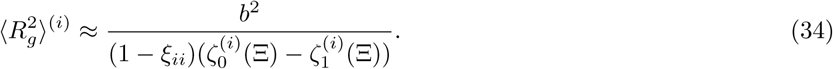

#### D. Encounter probability of monomers of the heterogeneous RCL polymer

We now compute the encounter probability (EP) between any two monomers of the heterogeneous RCL polymer, by first distinguishing between two cases: encounters between monomers of the same TAD (intra TAD), and monomers of different TADs (inter TAD).

##### 1. Intra-TAD encounter probabilities

We compute now the variance 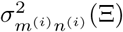 of the vectors between monomers 1 ≤ *m, n* ≤ *N*_*i*_ of TAD_*i*_ using normal coordinates 13

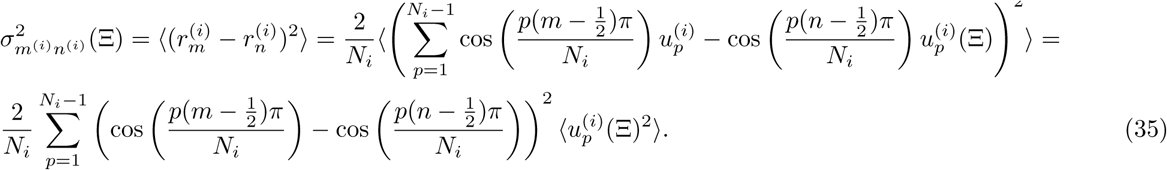

We recall that the variance of the normal coordinates in Eq. 21 is

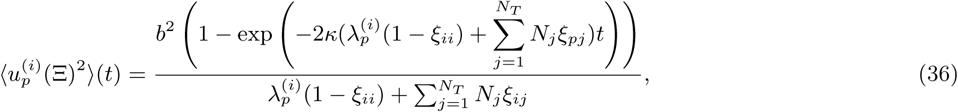

where 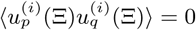, *p* ≠ *q*. The steady-state limit of 36 gives

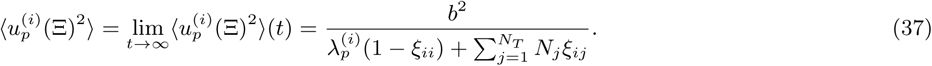

Substituting 37 in 35, we obtain

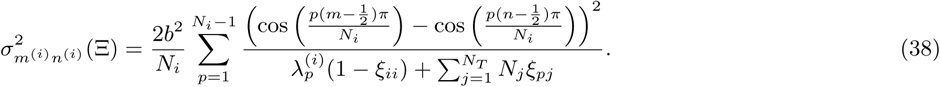

To compute the sum 38 for *N*_*i*_ » 1, we use the Euler-MacLaurin formula with the change of variable 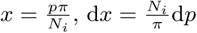

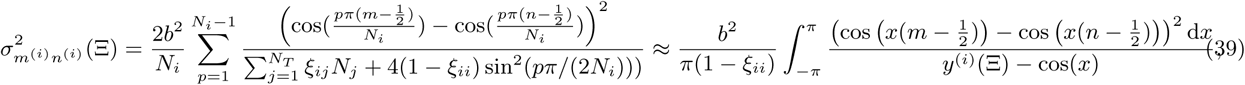

where *y*^(*i*)^(Ξ) is defined in expression 32. To compute the integral in 39 we make the change of variable *z* = exp(i_c_*x*), with i_c_ the complex unit, to obtain the expression

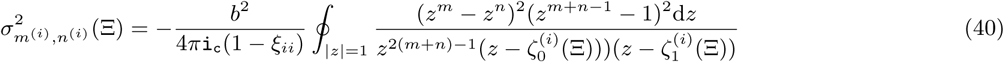

where 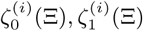 are defined in 33. The integrand in 40 has a pole of order 2(*m* + *n*) − 1 at *z* = 0 and a simple pole at 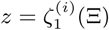. Since 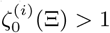 for all Ξ (Eqs. 32–33), this pole is outside the unit disk and does not contribute to integral. The final result is (see also [7] for details)

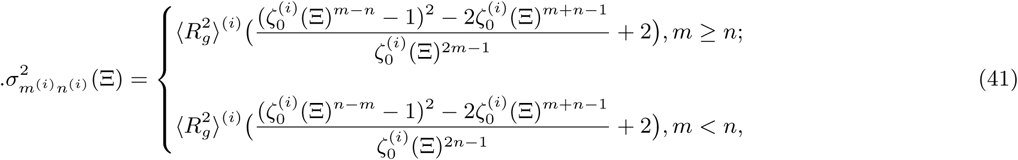

where 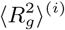 is the MSRG of TAD_*i*_ defined by expression 34. Because the heterogeneous RCL model belongs to the class of Gaussian models [12], the EP between monomer *m* and *n* of TAD_*i*_ is given by

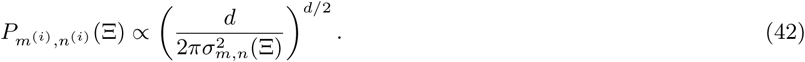

##### 2. Inter-TAD encounter probabilities

We now compute the EP between monomers of TAD_*i*_ and TAD_*j*_ (*i* ≠ *j*). We start by computing the variance of the vector between 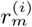 of TAD_*i*_ and 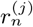 of TAD_*j*_ by using the position of center of masses 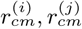, respectively:

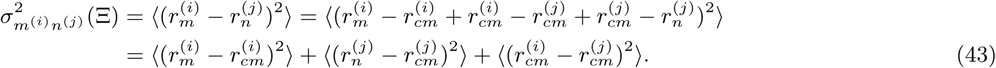

The variance of the vector between a monomer *m* of TAD_*i*_ and its center of mass in normal coordinates (Eq. 13) at steady-state is

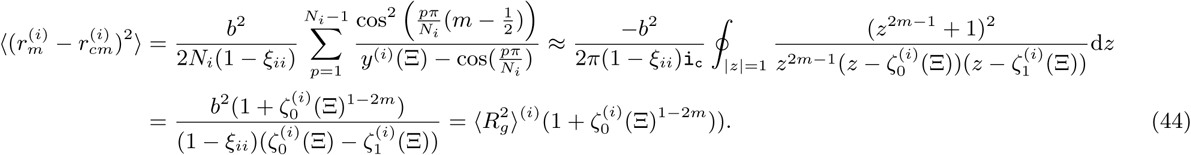

Similarly for TAD_*j*_, we obtain

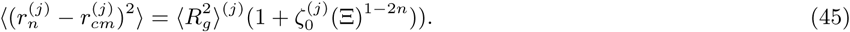

Under the assumption of weak inter-TAD connectivity (assumption *H*^1^, Eq. 29), the position of the centers of massed of TADs are independent. Thus, we obtain

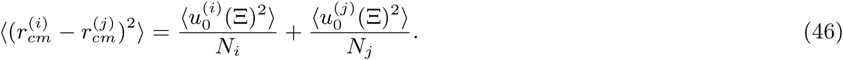

We compute the variance of the center of mass 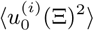 from equation 22 at steady-state, to obtain

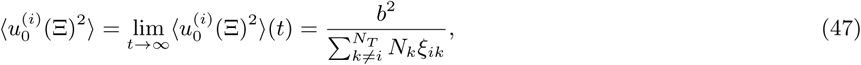

and similarly for the center of mass of TAD_*j*_

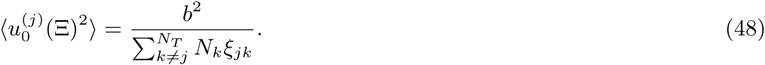

Substituting 48 and 47 into 46, we obtain the variance of the vector between centers of masses

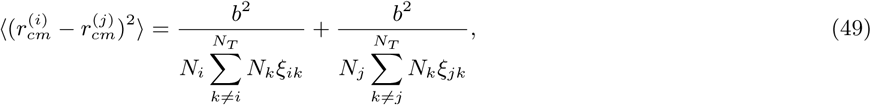

where here we impose a non-vanishing connectivity condition

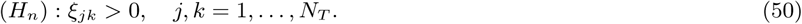

To obtain the final expression for the inter-TAD variance, we substitute expressions 44, 45 and 49 into 43, to have

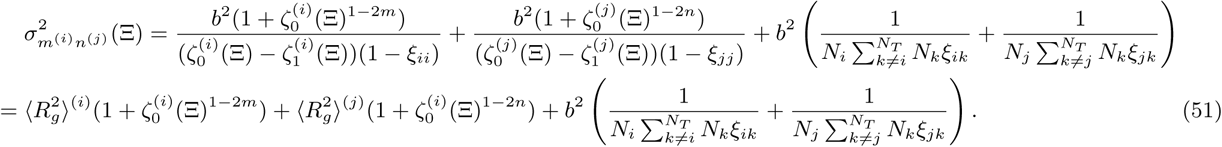

The EP between monomer m of TAD_*i*_ and *n* of TAD_*j*_ is therefore

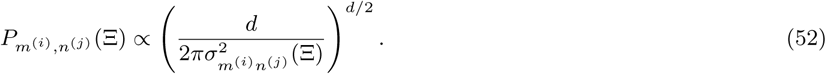

#### E. Mean-square displacement of monomers of the heterogeneous RCL polymer

We now derive an expression for the mean square displacement (MSD) of monomers in TAD_*i*_. The procedure follows the steps presented in [7], however we replace the mean number *Nξ* of connectors of each monomer of a single TAD by the the mean number 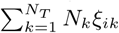 of connected monomers in TAD_*i*_ in a multi-TAD RCL. When the dominant TAD connectivity assumption *H*_1_ (Eq. 29) is satisfied, the relaxation times *τ*_p_(*ξ*), *p* = 1, ..*N*^(*i*)^ − 1 of the internal modes of the RCL polymer of *N*_*i*_ monomers, is [7]

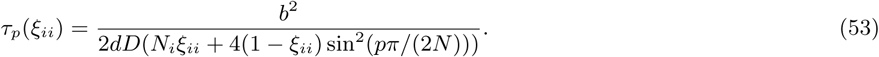

Thus, for intermediate time scales *τ*_*N*−1_(*ξ*_*ii*_) ≤ *t* ≤ *τ*_1_(*ξ*_*ii*_) the average MSD (over monomers and realizations *𝒢*) is

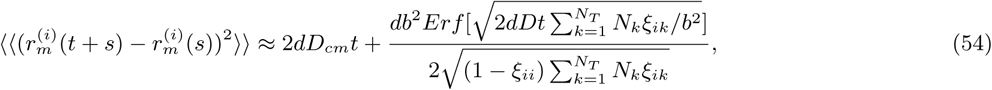

where 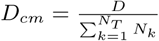. Expression 54 can further be approximated by

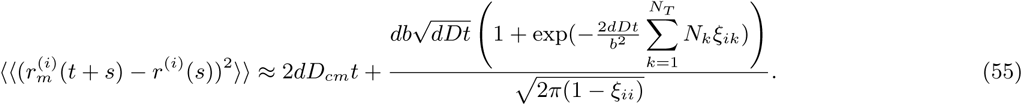

Expression 55 scales as 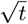 similar to the Rouse polymer. This effect is due to the averaging over all realizations. We also highlighted here how the MSD curve depends on the connectivity between TAD_*i*_ and all other TADs.

#### F. Distribution of the distance between monomers of the RCL polymer

We now derive an expression for the distribution of the distance between any two monomers *r*_*m*_ and *r*_*n*_ of the heterogeneous RCL polymer model. Since the RCL polymer belongs to the Gaussian chain family [12], the vector *r*_*m*_ − *r*_*n*_ is actually Normally distributed in any dimension *j* = 1, …, *d* with mean *µ*_*mn*_ = 0 and standard-deviation *σ*_*mn*_(Ξ) (square root of Eqs. 41, 51), where Ξ is the connectivity fraction matrix (Eq. 4, main text). The distance *D*_*mn*_ between monomers *r*_*m*_ and *r*_*n*_ is defined by

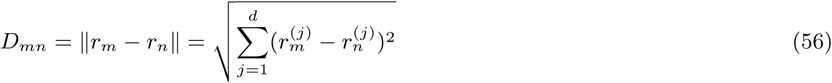

and is a *χ*-distributed random variable with *d* degrees of freedom, as the norm of *d*-dimensional Normally distributed vector *r*_*m*_ − *r*_*n*_. The normalized norm to the standard deviation

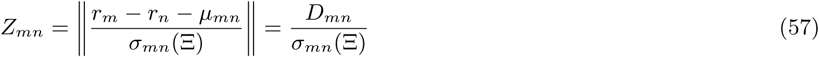

is distributed according to standard *χ* distribution, as the norm of d-dimensional Normally distributed random vector with mean zero and unitary standard-deviation. Thus, the distribution *f*_*D*_*mn*__(*x*) of the distances *D*_*mn*_ can be computed from the standard *χ* distribution

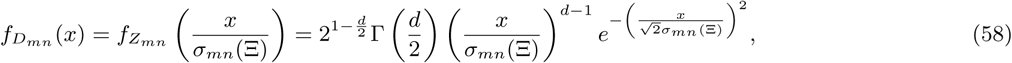

where Γ is the Γ−function.

Finally, using the average of *χ−* distribution 〈*Z*_*mn*_〉, we obtain for the average spatial distance between monomers *r*_*m*_ and *r*_*n*_

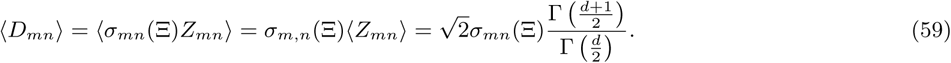

**TABLE I.** Mapping seven DNA FISH probe ends [18] at 3kb resolution from bp to monomers of the coarse-grained, three TAD, RCL polymer model with *N* = 605 monomers.

| Probe | Length (bp) | Monomer start ($r_m$) | Monomer end ($r_n$) |
| --- | --- | --- | --- |
| pEN1 | 9839 | 43 | 46 |
| pEN2 | 9612 | 109 | 112 |
| pLG1 | 9430 | 54 | 57 |
| pLG10 | 4503 | 116 | 117 |
| pLG11 | 4938 | 96 | 98 |
| X3 | 16060 | 73 | 78 |
| X4 | 12606 | 87 | 92 |

### III. COMPARISON OF THE DISTRIBUTION OF DISTANCES OF THE HETEROGENEOUS RCL MODEL WITH DNA FISH DATA

We compare the prediction of the distribution of three-dimensional distances between monomers of the heterogeneous RCL polymer to measurements of DNA FISH probe pairs. For this comparison, we use seven DNA FISH probe pairs of lengths 4-16kb, reported in [18]. We first binned the 5C-data [3] of TAD D,E, and F, at a monomer resolution of 3kb avoiding the two ends of a probe within the same monomer. We obtained a coarse-grained genomic section of *N* = 605 monomers, with *N*_*D*_ = 123, *N*_*E*_ = 174, *N*_*F*_ = 308. We then mapped the bp position of probes to monomers (see SI of [18]). In Table I we summarize the details of mapping FISH probes at 3kb to monomers.

We then constructed a three TAD RCL polymer of *N* = 605 monomers. To obtain the average connectivity fraction matrix Ξ, we fit the EP of the RCL polymer (Eqs. 42–52) to the empirical 5C EP data of TAD D, E, and F (see Methods section, main text) and obtained:

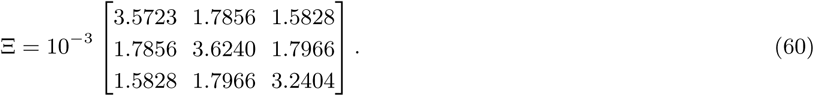

To compute the distribution of the three-dimensional distances *f*_*Dmn*_(*x*) (Eq. 58), we first compute the variance 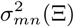 (Eqs. 41–51) of the distance between probes (monomers *m, n* Table I), where we substitute the fitted Ξ (Eq. 60) in Eq. 58, with *b* = 0.2*µm*. In Fig. 2, we plotted the distribution of 3D DNA FISH probe distances (black) vs. the predicted distribution *f*_*D*__*mn*_(*x*) (red), and further plotted the mean DNA FISH (green circles) and the predicted distances 〈*D*_*mn*_〉 (Eq. 59) of the RCL model, showing a very good agreement and confirming the predictive value of the RCL-polymer model.

### IV. HETEROGENEOUS RCL MODEL WITH VOLUME EXCLUSION FORCESHETEROGENEOUS RCL MODEL WITH VOLUME EXCLUSION FORCES

To examine the consequences of volume exclusion forces on the statistical properties of the heterogeneous RCL polymer (Eqs. 34, 42, 52), we now add to the potential in Eq. 1 the harmonic volume exclusion potential

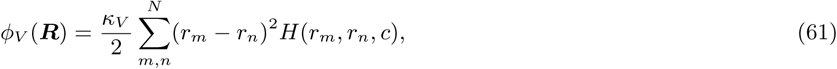

where *κ*_*V*_ is the spring constant, and *H*(*r*_*m*_, *r*_*n*_, *c*) is an indicator function for the distance between monomers *r*_*m*_ and *r*_*n*_ within a ball of radius *c*, defined as

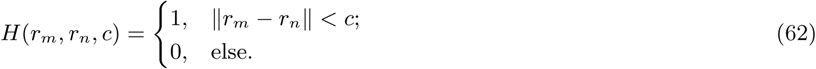

The exclusion radius *c* characterizes physical properties of external forces applied on the chromatin at a given coarse-grained scale. This parameter is thus independent of the mean distance *b* between connected monomers at that scale (see Method section, main text).

The total potential of the heterogeneous RCL polymer becomes

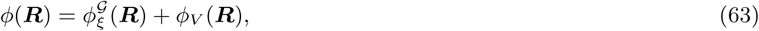

where 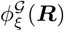 is the potential derived in Eq. 1. In the present model, we do not account for any topological constraints on the chromatin at any resolution. The result of this procedure is a heterogeneous RCL polymer with three TADs, similar to the one presented in Fig. 2 (main text). We performed 10,000 simulations with a time step ∆*t* = 0.01*s*, diffusion coefficient

**FIG.S 2.**
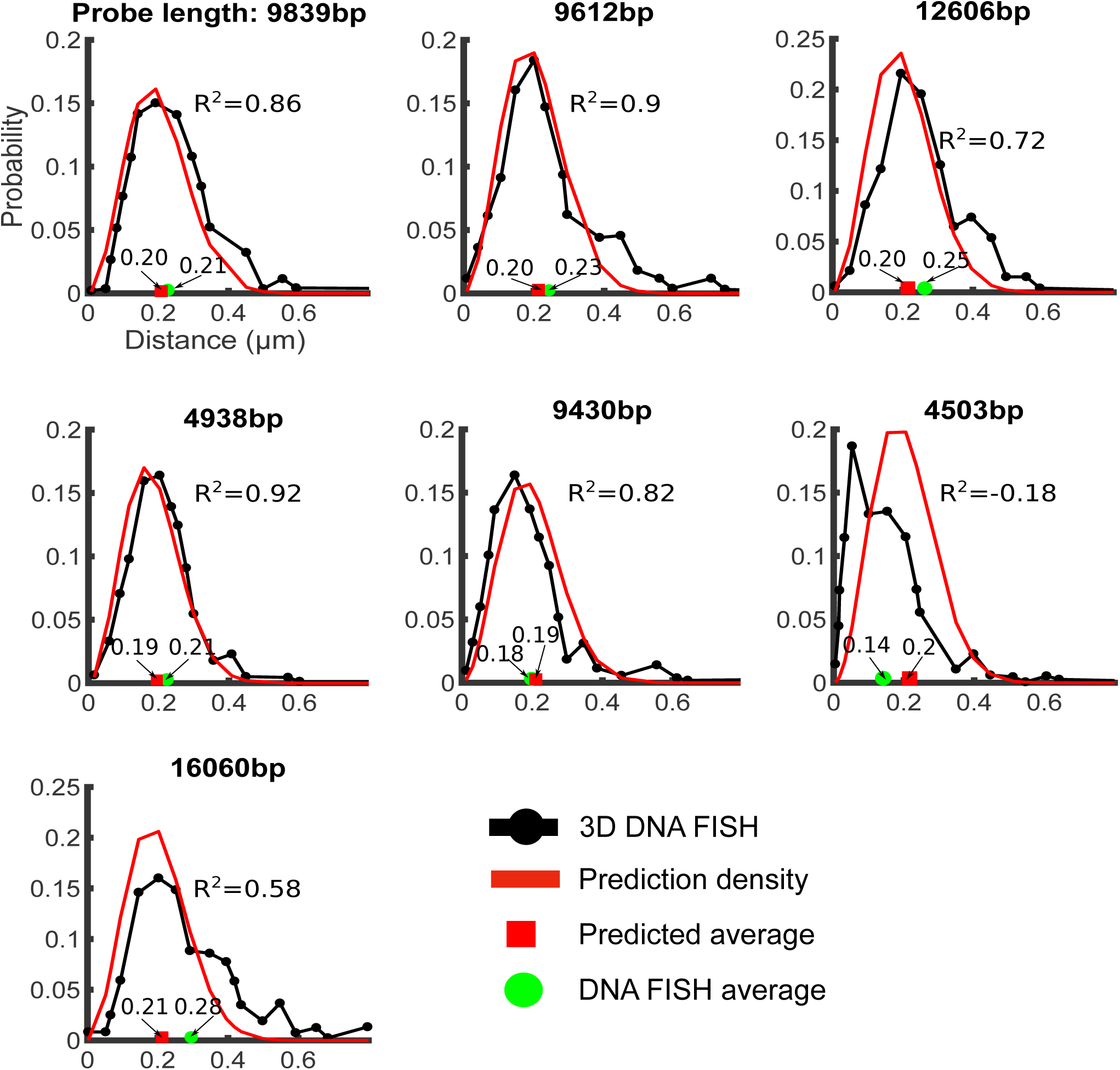
Comparing 3D DNA FISH with RCL model prediction. Distribution of three-dimensional distances between seven DNA FISH probe pairs (black) of lengths 4.5kb-16kb (measured in [18]) and predicted distributions *f*_*Dmn*_ (Eq. 58) of the RCL polymer model at 3kb resolution. Average values for DNA FISH (green circle) and predicted (Eq. 59, red squares).

*D* = 8 × 10^−3^ *µm*^2^*/s*, and the mean length *b* = 0.2*µm*. We examine two values of the radius of exclusion *c* = *b/*5*, b/*3*µm*, with an encounter distance *ε* = *b/*4 and *ε*= *b/*5*µm*, respectively.

In SI Fig. 3A, we plotted the encounter frequency matrix obtained from numerical simulations of Eq. 7 (main text) with the added volume exclusion potential 63 and a radius of exclusion *c* = 0.04*µm*, where the number of connectors within and between TAD satisfies the dominant intra-TAD connectivity assumption (Eq. 29). We find three distinct TADs and higher-order structures (meta-TADs) in the encounter frequency matrix (SI Fig. 3A), although the encounter frequency is reduced in comparison to 2A (main text), due to volume exclusion forces.

Interestingly, we find a good agreement between the EP obtained from numerical simulations and Eq. 42, 52, within and between TADs. In SI Fig. 3B, we plotted the EP for monomers *r*_20_, *r*_70_, and *r*_120_, located at the center of each TAD, against the EP expression Eqs. 42–52, and found a good agreement. The average MSD of monomers in each TAD (SI Fig. 3C) also confirm Eq. 54 for TAD_1_ (blue) and TAD_2_ (yellow) for all 0 *< t <* 25 s, and for TAD_2_(red) for all times *t* > 5 s. Finally, the mean radius of gyration obtained by simulations was 0.179, 0.134, and 0.168*µm*, which should be compared to 0.178, 0.132, and 0.167*µm* obtained from Eq. 54, with an average error of 1%. These results are remarkable and already show that the RCL polymer model is able to account to volume exclusion. We shall offer a possible explanation below.

**FIG.S 3.**
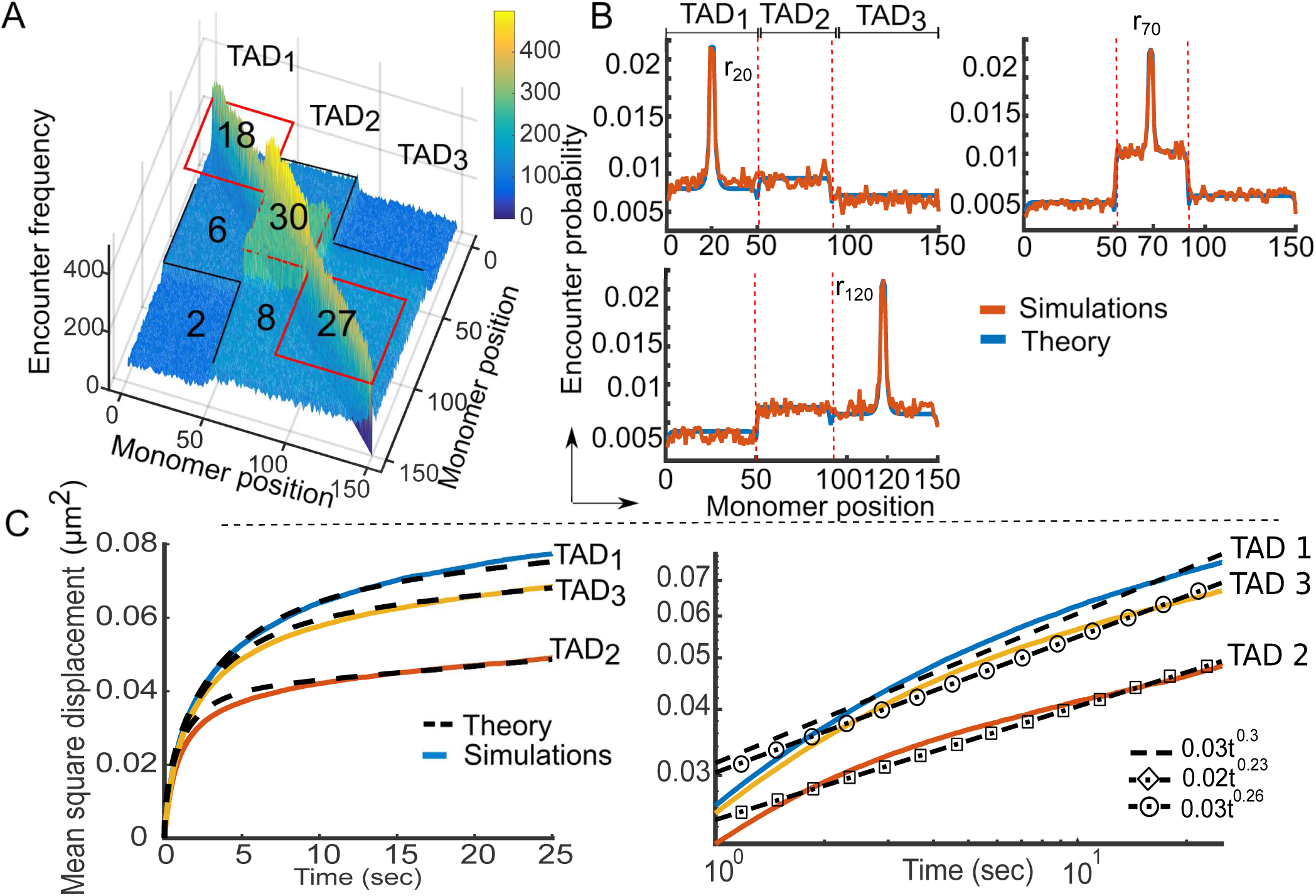
Statistical properties of the heterogeneous RCL polymer with added volume exclusion. **A.** Encounter frequency matrix of a polymer with three TAD blocks (TAD_1_, TAD_2_, TAD_3_) of *N*_1_ = 50, *N*_2_ = 40, *N*_3_ = 60 monomers is computed from 10,000 simulations of the RCL polymer using Eq. 7 (main text) with the added volume exclusion potential 63 and ∆*t* = 0.01 *s*, *D* = 8 × 10^−3^ *µm*^2^*/s*, *d* = 3, *b* = 0.2*µm*, *ε* = 0.02*µm*, and *c* = 0.04*µm*. The number of added connectors within and between TADs appears in each block. Three distinct diagonal TADs are visible (red boxes) where secondary structure appears (black lines) due to weak inter-TAD connectivity. **B.** Encounter probability (EP) of the heterogeneous RCL described in panel A, where the simulated EP (red) agrees with the theoretical one (blue, Eqs. 42, 52). We plot the EP of the middle monomer in each TAD: monomer *r*_20_ (Upper left), monomer *r*_70_ (Upper right) and monomer *r*_120_ (bottom left), where TAD boundaries are in vertical dashed red lines. **C.** Averaged mean squared displacement (left panel) of monomers in TAD_1_ (blue), TAD_2_ (orange) and TAD_3_ (yellow), using simulations of RCL polymer, as described in panel A: simulation (continuous line) vs. theory (dashed, Eq. 54). (Right)MSD plot in log-log scale. The estimated anomalous exponents uses the least square method in the log space of the a model 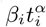 (Eq. 17, main text). For TADs *i* = 1..3 in the intermediate time of 1 to 25s results in *α*_1_ = 0.3, *α*_2_ = 0.23, *α*_3_ = 0.26 for TAD_1_, TAD_2_, and TAD_3_, respectively, showing how a given connector configuration defines the anomalous exponents.

We then repeated the same protocol after increasing the exclusion radius to *c* = 0.067*µm* (SI Fig. 4) and found that the encounter frequency (panel A) still contains three distinct TADs with meta-TAD like higher organization, but some discrepancies appeared between the EP expression (SI Fig. 4B, blue, Eq. 42, 52) and the simulation EP (orange). We further found disagreements between MSD from simulations (SI Fig. 4C, continuous line) and expression 54 (dashed). Finally, the mean radius of gyration from simulations is 0.192, 0.145, 0.181*µm* vs. 0.178, 0.132, and 0.167*µm* obtained from Eq. 34, for TADs 1 to 3, respectively, an average error of 8.5%.

We conclude from these results that the steady-state and dynamic properties (Eqs. 34, 54) are adequate to describe the chromatin using heterogeneous RCL polymer with volume exclusion radius of up to 40 nm. Possibly, many long-range forces are responsible to generate a polymer statistics that contain the one generated with volume exclusion. This result can be due to the homogenous distribution of connectors that on average pushes away all monomers.

**FIG.S 4.**
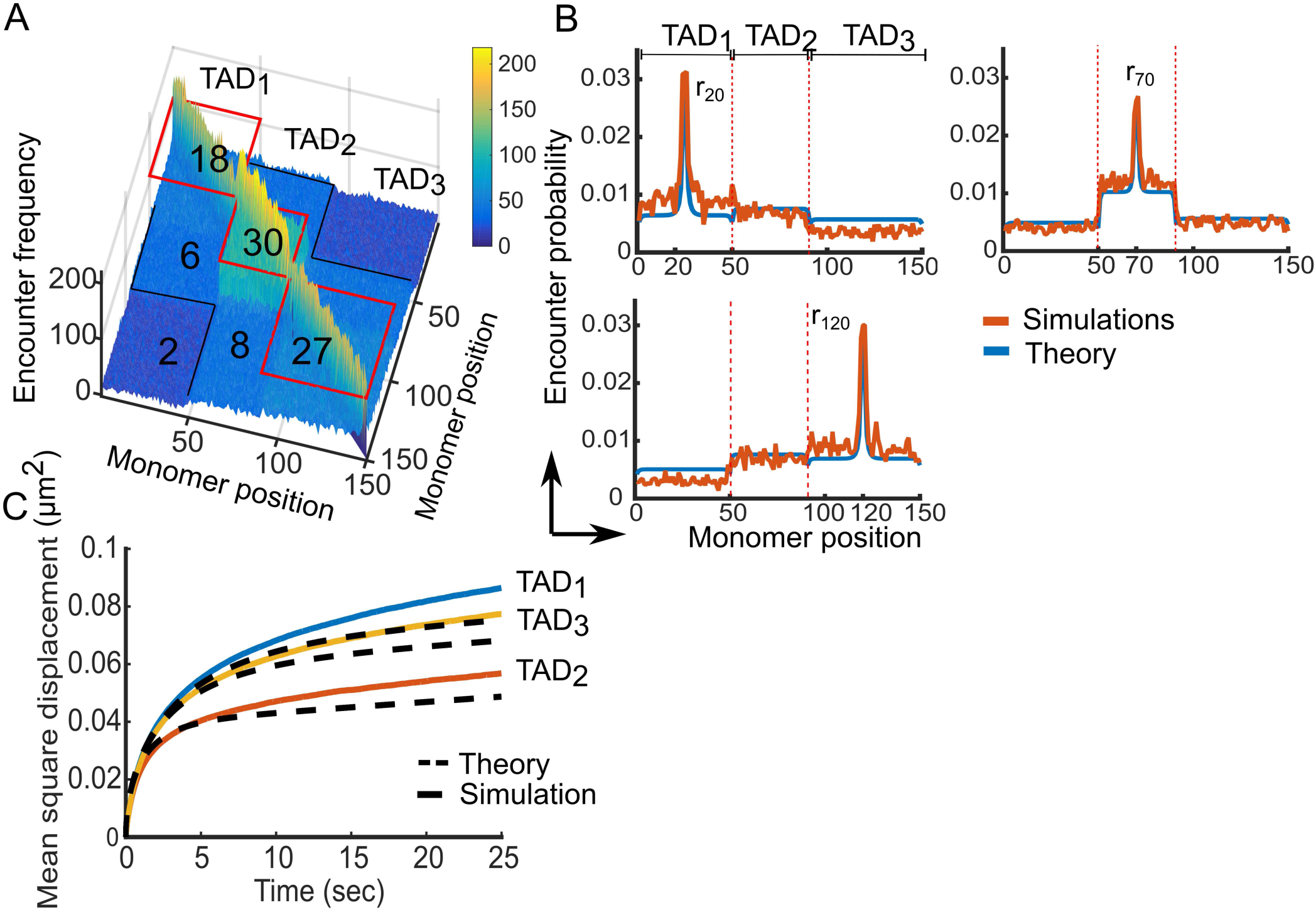
Statistical properties of the heterogeneous RCL polymer with added volume exclusion. **A.** Encounter frequency matrix of a polymer as in SI Fig. 3A, with a radius of exclusion *c* = 0.067*µm*. The number of added connectors appears in each block. Three distinct diagonal TADs are visible (red boxes) where secondary structure appears (black lines) due to weak inter-TAD connectivity. **B.** Encounter probability (EP) of the heterogeneous RCL described in panel A, where the simulation EP (orange) deviates from the theoretical EP (blue, Eqs. 42, 52), plotted for the middle monomer in each TAD: monomer *r*_20_ (Upper left), monomer *r*_70_ (Upper right) and monomer *r*_120_ (bottom left), and TAD boundaries appear in vertical dashed lines. **C.** Averaged mean squared displacement of monomers in TAD_1_ (blue), TAD_2_ (red), and TAD_3_ (yellow), using simulations of RCL polymers, as described in panel A: simulations (continuous line) vs. theory (dashed, Eq. 54), that do not agree for *t* > 5s.

### V. COMPARING RCL POLYMERS RECONSTRUCTED FROM TWO REPLICAS OF 5C DATA AT 10KB RESOLUTION

To test the validity of the present approach in reconstructing the statistics of several interacting TADs, we use each replica of the 5C data of the X chromosome [3], TAD D, E, and F, to find and compare the average number of connectors between replicas. We started by fitting the steady-state EP (Eqs. 42 and 52) to each replica independently. We then bin the 5C data of TADs D, E, and F [3], at 10kb resolution (Methods, main text and also [6]), which resulted in *N*_*T*_ = 183 monomers, where *N*_*D*_ = 37, *N*_*E*_ = 53, and *N*_*F*_ = 93, respectively. In Fig. 5A, we plotted the average number of connectors found by fitting the EP of three TADs across three stages of cell differentiation: mESC, NPC, and MEF, for replica 1 (SI Fig. 5A) and replica 2 (SI Fig. 5B), where we approximate the number of connectors to the nearest integer. In SI Fig. 5C, we plotted the absolute difference |*Nc*(*Rep*1) − *Nc*(*Rep*2)|, between the number of connectors *Nc*(*Rep*1) (resp. *Nc*(*Rep*2)), found for replica 1 (resp.2). The RCL fit to empirical data resulted in a small difference (0-2 connectors) between the two replicas, where the only significant difference of 5 connectors was found for intra-TAD connectivity of TAD F (Fig. 5C, left).

To examine the robustness of the RCL fitting procedure, we tested the fit of the EP of one replica of the 5C data to the empirical EP of the second replica, on the basis of individual monomers. We first fitted the empirical EP of each monomer of replica 1 for TADs D, E, and F, at a resolution of 10kb, by performing 183 curve fitting using Eqs. 42, 52 to obtain the connectivity fractions *ξ*_*m*_, *m* = 1, …, 183 of each monomer. Using the fitted *ξ*_*m*_ from replica 1, we computed the EP 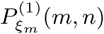, *n* = 1, …, 183, between monomers *m* and *n* of replica 1 and compared it to the empirical EP *E*^(2)^(*m, n*) of replicas 2 using the norm

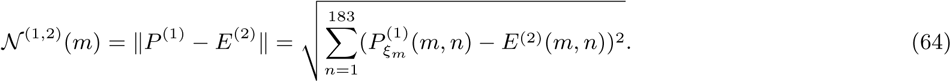

We repeated the fitting process described above when we exchanged the roles of replica 1 and 2, and computed the norm 𝒩^(2,1)^(*m*) (Eq. 64) for each monomer *m* = 1*, …,* 183. In Fig. 5D, we plotted the norm 64 when we fitted the EP of replica 1 and compared it with the empirical EP of replica 2 (left), and when we fitted replica 2 and compared it with the empirical EP of replica 1 (right). We found that for a majority of monomers the norm satisfies 0.1 ≤ 𝒩 (*m*, 1, 2) ≤ 0.3, for each 𝒩 (*m*, 1, 2) (left) and *N*(*m,* 2, 1) (right). The peaks at *m* = 57, 160 indicate disagreement between fitted EP of replicas 1 and 2 with empirical EP of replica 2 and 1, respectively, due to the presence of persistent long-range looping (peaks of the empirical EP matrix) for these monomers. We further computed the average of the norm 64 over all monomers *m* = 1, …, 183 by

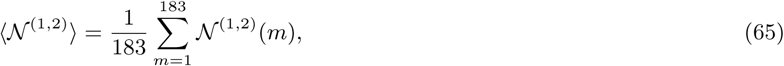

and obtained 〈𝒩^(1,2)^〉 = 0.12, 0.11, and 0.12, for mESC, NPCs and MEF, respectively. Reversing the roles of replica 1 and 2, we obtained 〈𝒩^(2,1)^〉 = 0.13, 0.1, and 0.13, for mESC, NPCs, and MEF, respectively. From the results in SI Fig. 5, we concluded that the present fitting procedure using RCL polymer is consistent across replicas of the 5C data. We then compared the mean radius of gyration (MRG) and compaction ratio of each 5C replicas in stages: mESC, NPC, and MEF, (Fig. 6). To compute the MRG, we first computed the value of *b*, the mean square distance between connected monomers, at a scale of 10kb. For this end, we used the value of *b* = 0.2*µm* at 6kb resolution (Figs.2–3, main text), and chose the criteria that the MSRG (Eq. 34) must be invariant across scales in order to compute the value of *b* at 10 kb. Therefore, we obtain the equation for TAD_*i*_ for 6kb and 10 kb, we have

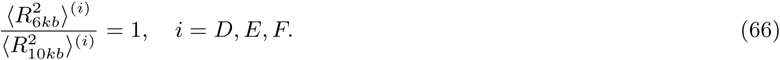

Using Eq. 34, we obtain

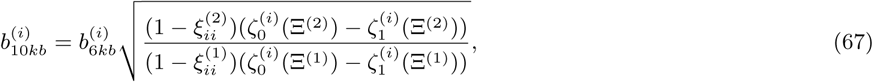

where 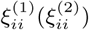, Ξ^(1)^, (Ξ^(2)^) are found by fitting the EP (Eq. 42, 52) to the empirical data at 6kb (10kb). Applying Eq. 67 to the 5C data at 6 and 10 kb resolution, mESC, for TADs D, E, and F, we obtain the value of *b*_10*kb*_ for each TAD: *b*_*D*_ = 0.185, *b*_*E*_ = 0.16, *b*_*F*_ = 0.2. In simulations, we use a single *b* value, obtained by averaging *b*_*D*_, *b*_*E*_, and *b*_*F*_ to obtain *b*_10*kb*_ = 0.1814*µm*.

We found that the average (over TADs) of the MRG of the 5C replica 1 is 0.2*µm* at mESC stage and dropped to 0.186 *µ* at NPC stage, but has an average of 0.2*µm* at MEF stage (SI Fig. 6A, left). These compaction and decompaction changes of TADs across differentiation are present for 6kb resolution (Fig. 3C, main text). The compaction ratio (Eq. 16, main text) of replica 1 (SI Fig. 6A, right) increased from an average value of 19 at mESC stage to 28 at NPC stage, and was 20 at MEF stage. The compaction ratio for the 6 TADs is qualitatively in agreement those for the average of the two replicas for three TADs (Fig. 3C, main text) across cellular differentiation. However, we found several differences from the synchronous compaction and decompaction of TADs in the MRG of replica 2 (SI Fig. 6B, left) where we observed a slow monotonic decrease in the MRG from an average of 0.21 *µ* at mESC stage to 0.2*µm* at MEF stage.

### VI. SENSITIVITY OF THE RCL POLYMER MODEL TO TAD BOUNDARIES’ LOCATIONS

To examine the robustness of the RCL model, we varied the number of TADs and the position of their boundaries. For this end, we use the 5C data in Fig. 3 (main text) at 6kb and further subdivide each TAD, D, E, and F in half. After subdivision we obtain six sub-TADs with *N*_1_ = 31, *N*_2_ = 31, *N*_3_ = 43, *N*_4_ = 44, *N*_5_ = 77, *N*_6_ = 77. We then repeated the procedure of EP fitting (Method section) for three stages of cell differentiation, to obtain the average number of added connectors within and between TADs (SI Fig. 7A). To compare the number of connectors of the three TAD case to those with no subdivision (Fig. 3C, main text), we average each non-overlapping 2 × 2 blocks of (SI Fig. 7A), to obtain the expected number of monomers in a 3 × 3 matrix corresponding to TADs D, E, and F (SI Fig. 7B). We then computed the absolute difference between the 2 × 2 block average associated with each TAD in SI Fig. 7B to the results obtained for 3 TADs (Panel C). We find differences of 0-30% connectors in all three cell types, with the most significant being six connectors in the intra-TAD connectivity of TAD E in the MEF stage. The difference between the number of connectors within TADs is 4, 4.5, 4, for mESC, NPC, and MEF, respectively, whereas the difference of inter-TAD connectors is 2, 0.7, and 1.7, for mESC, NPC, and MEF, respectively. We conclude that the subdivision of TADs and position of TAD boundaries across experiments can lead to differences in the intra-TAD connectivity of up to 30%.

**FIG.S 5.**
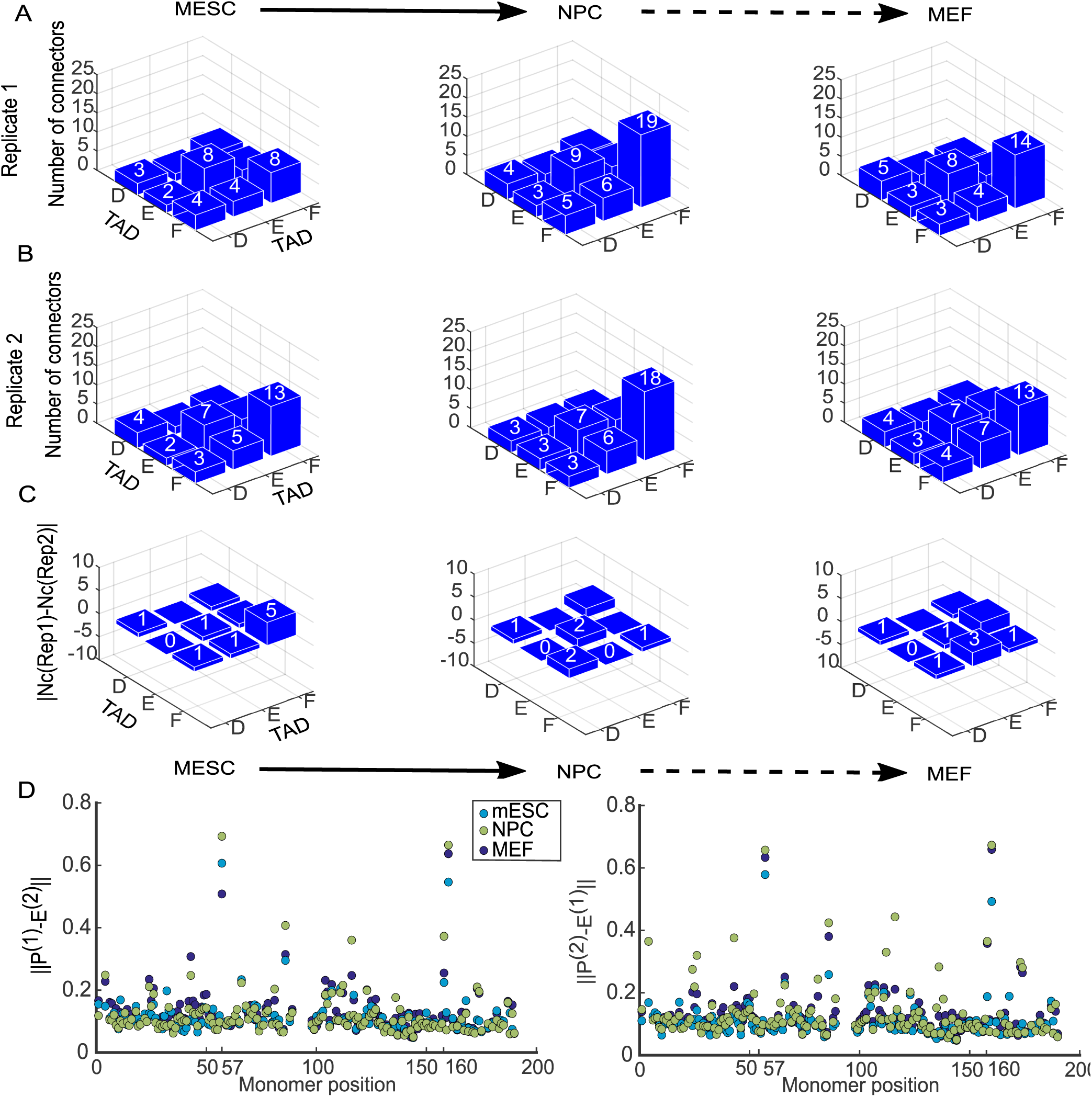
Comparing the number of connectors between two replica of 5C data at a resolution of 10kb. **A.** Average numbers of inter and intra-TAD connectors for TAD D, E, and F, obtained by fitting the RCL model EP (Eq. 42, 52) to each monomer of replica 1 of the dataset presented in [3], binned at 10 kb resolution, and for three stages of cell differentiation: mESC (left), NPC (middle), and MEF (right). **B.** Same as in panel A for replica 2. **C.** Absolute difference between the average number of connectors found independently for replica 1 and 2 (panels A and B), and for the three cell stages. **D.** Norm of the difference ||*P*^(1)^ *− E*^(2)^|| (Eq. 64), between the EP *P*^(1)^ fitted to replica 1, and the empirical EP *E*^(2)^ of replica 2 (left) and then we show the opposite (right), where we compute the norm of the difference ||*P*^(2)^ *− E*^(1)^||.

**FIG.S 6.**
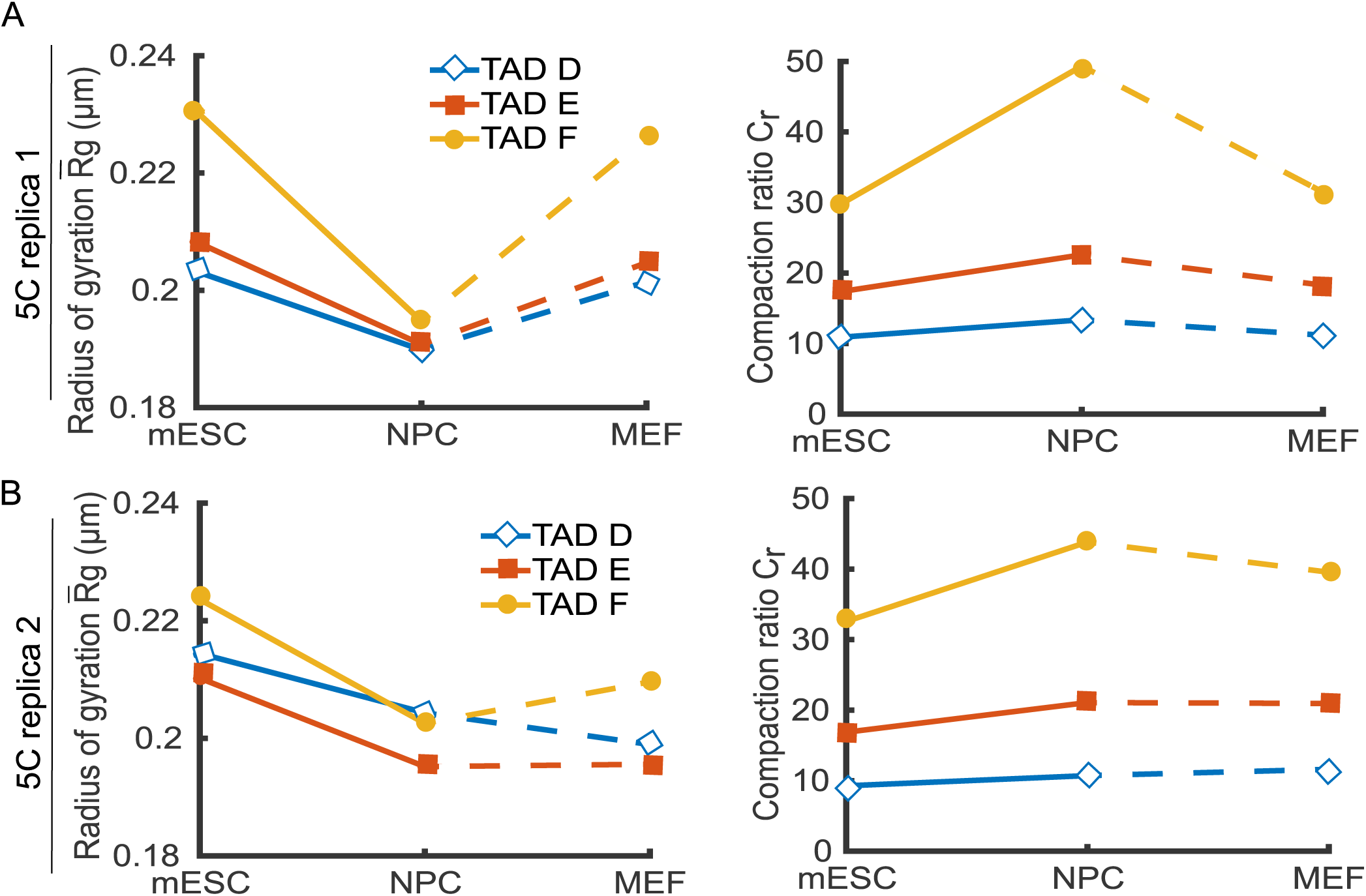
Statistics of individual replicas of the 5C data. **A.** Mean Radius of Gyration (MRG) of replica 1 of the X chromosome (left) at 10 kb resolution for TADs D (blue diamonds), E (orange squares), and F (yellow circles), and for three stages of cell differentiation, shows a synchronous TAD compaction in the transition from mESC to NPC cells, and the compaction ratio at the MEF stage. The compaction ratio (right, Eq. 16, main text) further reveals the chromatin changes across cell differentiation. **B.** Mean radius of gyration (left) of TADs D (blue diamonds), E (red, squares), and F (yellow, circles) of 5C replica 2 of the X chromosome indicates a synchronous compaction of all three TADs in the transition from mESC to NPC. The changes in the compaction ratio (right) of replica 2 is comparable to that of replica 1 (panel A, right).

### VII. CHROMATIN DYNAMICS THROUGHOUT DIFFERENTIATION FOR 6 TADS

We study the consequence of dividing the three TADs into six, by subdividing each in half (see previous subsection). We computed the mean radius of gyration for TADs 1-6 (SI Fig. 8A left), and obtained an average (over TADs) of 0.13*µm* at mESc stage, 0.12*µm* for NPC, and 0.13*µm* at MEF. This synchronous compaction and decompaction of the six TAD case at 6 kb resolution is in agreement with our finding for the three TAD case (SI Fig. 8A, right) for which we find that the mean radius of gyration decreased from 0.21*µm* at mESC stage, to 0.19*µm* at NPC, and was 0.2*µm* for MEF stage.

The mean radius of gyration for sub-TADs do not add up to that of the whole TAD and this implies that sub-TADs are intermingled. These configurations can further be accounted for by the difference in the number of added connectors between the six and three TAD case (SI Fig.7D). We further computed the compaction ratios (Eq. 16, main text) for the six TAD case (SI Fig. 8B, left), which increase from an average of 93 at mESC stage, to 115 at NPC and dropped back to 90 at MEF stage. The compaction ratio for the three TAD case (SI Fig.8, right), increased from 56 at mESC stage to 81 at NPC, and decreased to 62 at MEF stage.

We conclude from the results in Fig. 8 that the qualitative compaction and decompaction of the three TADs throughout differentiation is preserved between the three TAD and after subdivision into six TADs. However, we can expect a quantitative differences of up to 30% in the number of intra-TAD connectors.

**FIG.S 7.**
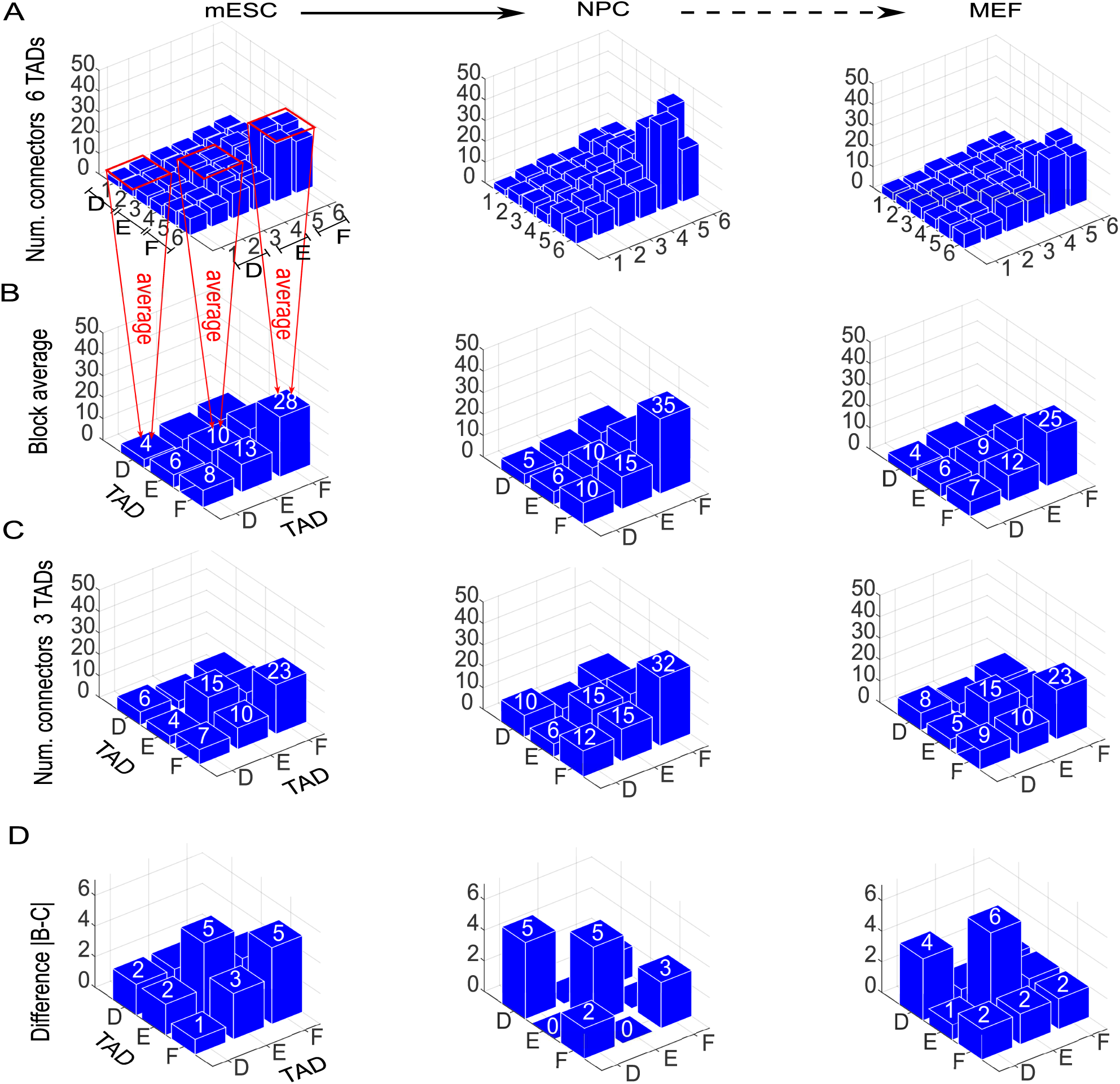
Robustness of TAD boundaries in RCL representation. **A.** Average number of connectors for the genomic section spanning TAD D, E, and F, when each TAD is further divided in half to give rise to six TADs, binned at 6kb resolution, for mESC (left), NPC (middle), and MEF (right) cell types. **B.** Average number of connectors obtained by fitting SI Eq. 42, 52 to the empirical EP, in each non-overlapping 2 × 2 blocks of the number of connectors in panel A, corresponding to the initial subdivision of the genomic segment into TADs D, E, and F. **C.** Average number of added connectors for TADs D, E, and F presented in Fig. 3 (main text). **D.** Difference between the average number of connectors shown in panel B and panel C.

### VIII. COMPARING HIC/5C DATA RECONSTRUCTED FROM RCL POLYMER

To test the robustness of the present approach, we decided to compare the statistics and genome reorganization of the 5C, that we previously analyzed with HiC data [10] for the X chromosome at 10kb, of a genomic section around TADs D, E, and F over three successive stages of cell differentiation: mouse embryonic stem cells (mESC), neuronal progenitors (NPC), and cortical neurons (CN). The data for mESCs, NPC and CN are obtained from the GEO repository GSE96107 [10]. We use HiC-Pro v2.9.0 [16] to map the raw data to mouse reference genome mm10 (using Bowtie2 v2.3.0; [17]) and we aligned reads, with default settings to remove duplicates, assign reads to DpnII restriction fragments and filter for valid interactions. We generated a binned interaction matrices at 10kb resolution from the valid interactions and we use the Iterative Correction and Eigenvector decomposition method (ICE) implemented in HiC-pro to normalize the results. Finally, we mapped the boundaries of TADs according to the TAD boundaries reported in the 5C data [3].

**FIG.S 8.**
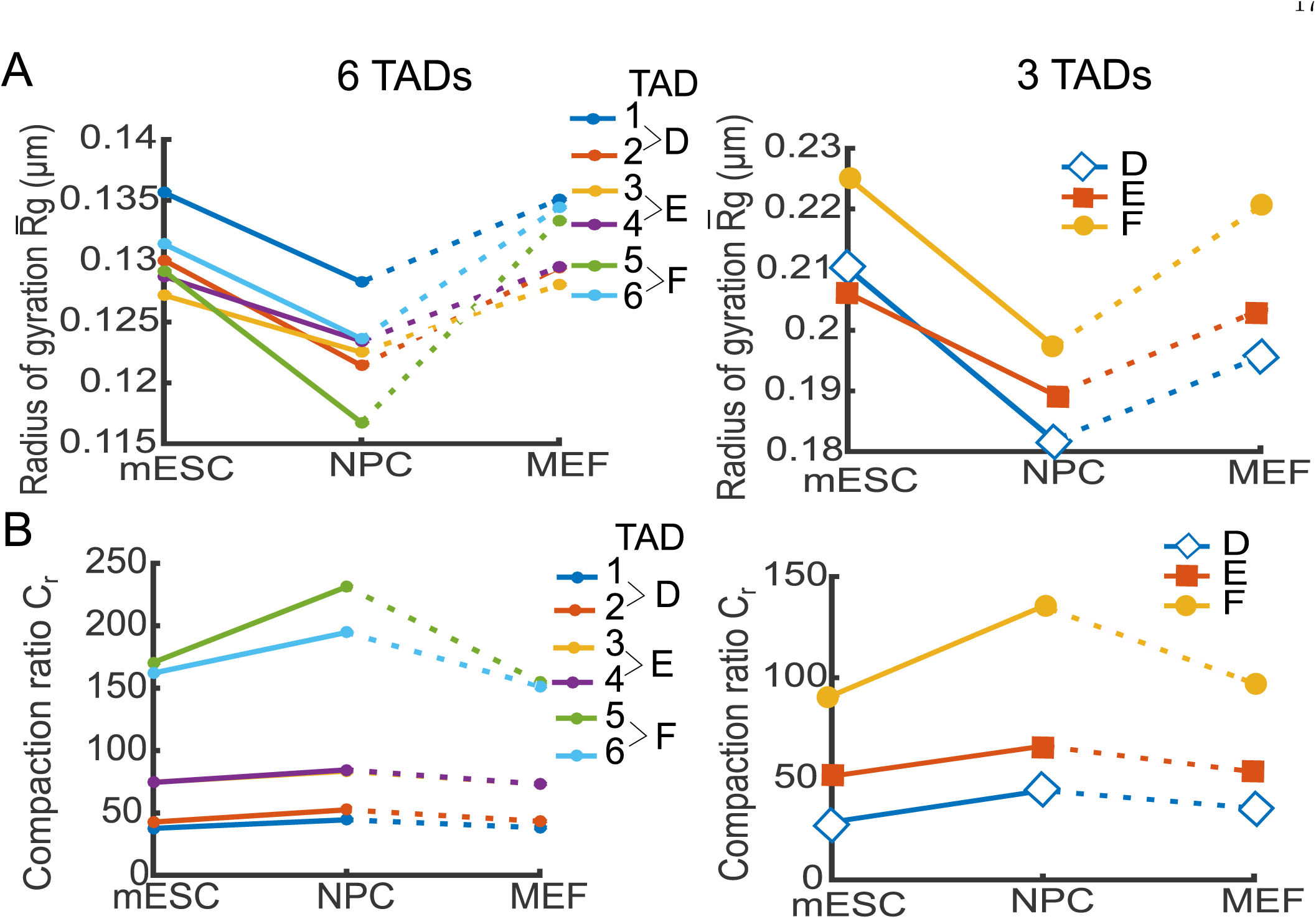
Comparing three and six sub-TADs throughout cellular differentiation. **A.** Mean radius of gyration of the 5C data [3] at 6 kb for TADs D, E, and F when each TAD is further divided in half (left), such that TADs D, E, and F comprise TADs 1-2, 3-4, and 5-6, respectively, and (right) with no TAD sub-division. **B.** Compaction ratio (left, Eq. 16, main text) of TADs 1-6 throughout differentiation, and with no TAD sub-division (right).

After binning at 10 kb, we obtained a coarse-grained genomic section of 183 monomers, where *N*_*D*_ = 37, *N*_*E*_ = 53, and *N*_*F*_ = 93, for TADs D, E, and F, respectively. We fitted the EP (Eq. 52 of 42) to the empirical EP of the coarse-grained HiC data to obtain the average number of connectors within and between TADs (Fig. 4A) for mESC (left), NPC (middle), and CN (right) cell types. We find a monotonic slow increase in the intra-TAD connectivity in the transition from mESC to CN cells, with very low inter-TAD connectivity (average 1.5 connectors) throughout all three stages of differentiation. The acquisition of connectors throughout differentiation affects the compaction of each TAD and is indicated by the mean radius of gyration (Fig. 4B), which steadily decreased from an average (over TADs) of 0.25 *µ* at mESC stage to 0.24*µm* at CN stage. The compaction ratio (Eq. 16, main text) slightly increases from an average of 10 at mESC stage to 14 at CN stage. We then compared the reconstructed HiC statistics with that of the 5C data [3] binned at 10 kb. We find that the average number of connectors of the 5C of TAD D and E (Fig. 4C) is in agreement with that of the HiC (Fig. 4A) for mESC and NPC cell stages. Cortical neurons cells are more advanced differentiation stage than the MEF and therefore the reconstructed statistics of the HiC CN and 5C MEF cannot be directly compared. The mean radius of gyration (Fig. 4D, left) for the 5C, remained at a average value of 0.2*µm* throughout the three staged of differentiation. However, the average compaction ratio (Fig. 4D, right), in mESC is 21.5, and increased to 24 at NPC and MEF stages. Taken together, we find consistency in the intra-TAD connectivity between the 5C and HiC data. However, the synchronous compaction of TADs at the differentiation from mESC to NPC is not seen for the HiC. This is in contrast to the TAD dynamics of the 5C for 10 kb resolution, the average of two replicas at 6kb, and for individual 5C replicas (Fig. 6). We attribute this discrepancy to the reduced inter-TAD connectivity of the HiC data in comparison to the 5C, and to the binning at 10 kb, which eliminated inter-TAD connectivity.

## References

[1] E. P. Nora, B. R. Lajoie, E. G. Schulz, L. Giorgetti, I. Okamoto, N. Servant, T. Piolot, N. L. van Berkum, J. Meisig, J. Sedat, et al., Nature 485, 381 (2012).

[2] M. Simonis, P. Klous, E. Splinter, Y. Moshkin, R. Willemsen, E. De Wit, B. Van Steensel, and W. De Laat, Nature genetics 38, 1348 (2006).

[3] B. D. Pope, T. Ryba, V. Dileep, F. Yue, W. Wu, O. Denas, D. L. Vera, Y. Wang, R. S. Hansen, T. K. Canfield, et al., Nature 515, 402 (2014).

[4] J. Dekker, K. Rippe, M. Dekker, and N. Kleckner, science 295, 1306 (2002).

[5] J. R. Dixon, S. Selvaraj, F. Yue, A. Kim, Y. Li, Y. Shen, M. Hu, J. S. Liu, and B. Ren, Nature 485, 376 (2012).

[6] Q. Szabo, D. Jost, J.-M. Chang, D. I. Cattoni, G. L. Papadopoulos, B. Bonev, T. Sexton, J. Gurgo, C. Jacquier, M. Nollmann, et al., Science Advances 4, eaar8082 (2018).

[7] M. Tark-Dame, H. Jerabek, E. M. Manders, D. W. Heer-mann, and R. van Driel, PLoS Computational Biology 10, e1003877 (2014).

[8] L. Giorgetti, R. Galupa, E. P. Nora, T. Piolot, F. Lam, J. Dekker, G. Tiana, and E. Heard, Cell 157, 950 (2014).

[9] O. Shukron and D. Holcman, PLOS Computational Biology 13, e1005469 (2017).

[10] E. Lieberman-Aiden, N. L. Van Berkum, L. Williams, M. Imakaev, T. Ragoczy, A. Telling, I. Amit, B. R. Lajoie, P. J. Sabo, M. O. Dorschner, et al., science 326, 289 (2009).

[11] O. Shukron and D. Holcman, Physical Review E 96, 012503 (2017).

[12] J. Fraser, C. Ferrai, A. M. Chiariello, M. Schueler, T. Rito, G. Laudanno, M. Barbieri, B. L. Moore, D. C. Kraemer, S. Aitken, et al., Molecular systems biology 11, 852 (2015).

[13] M. Doi and S. Edwards, The Theory of Polymer Dynamics Clarendon (Oxford, 1986).

[14] M. Bohn and D. W. Heermann, PloS One 5, e12218 (2010).

[15] I. Sokolov, Physical Review Letters 90, 080601 (2003).

[16] M. Bohn, D. W. Heermann, and R. van Driel, Physical Review E 76, 051805 (2007).

[17] D. W. Heermann, Current Opinion in Cell Biology 23, 332 (2011).

[18] J. S. Verdaasdonk, P. A. Vasquez, R. M. Barry, T. Barry, S. Goodwin, M. G. Forest, and K. Bloom, Molecular cell 52, 819 (2013).

[19] P. A. Vasquez and K. Bloom, Nucleus 5, 376 (2014), pMID: 25482191, https://doi.org/10.4161/nucl.36275.

[20] A. Amitai and D. Holcman, Physical Review E 88, 052604 (2013).

[21] J. Langowski and D. W. Heermann, in Seminars in cell & developmental biology, Vol. 18(5) (Elsevier, 2007) pp. 659–667.

[22] P. A. Vasquez, C. Hult, D. Adalsteinsson, J. Lawrimore, M. G. Forest, and K. Bloom, Nucleic Acids Research 44, 5540 (2016).

[23] S. Jespersen, I. Sokolov, and A. Blumen, The Journal of Chemical Physics 113, 7652 (2000).

[24] J. Bryngelson and D. Thirumalai, Physical review letters 76, 542 (1996).

[25] D. Jost, P. Carrivain, G. Cavalli, and C. Vaillant, Nucleic acids research 42, 9553 (2014).

[26] D. Michieletto, E. Orlandini, and D. Marenduzzo, Physical Review X 6, 041047 (2016).

[27] D. Noordermeer and W. de Laat, IUBMB life 60, 824 (2008).

[28] G. Fudenberg, M. Imakaev, C. Lu, A. Goloborodko, N. Abdennur, and L. A. Mirny, Cell reports 15, 2038 (2016).

[29] A. Amitai and D. Holcman, Physical Review Letters 110, 248105 (2013).

[30] T. M. Cheng, S. Heeger, R. A. Chaleil, N. Matthews, A. Stewart, J. Wright, C. Lim, P. A. Bates, and F. Uhlmann, Elife 4 (2015).

[31] D. E. Anderson, A. Losada, H. P. Erickson, and T. Hirano, The Journal of cell biology 156, 419 (2002).

[32] C. Weinreb and B. J. Raphael, Bioinformatics 32, 1601 (2015).

[33] S. S. Rao, M. H. Huntley, N. C. Durand, E. K. Stamenova, I. D. Bochkov, J. T. Robinson, A. L. Sanborn, I. Machol, A. D. Omer, E. S. Lander, et al., Cell 159, 1665 (2014).

[34] B. Bonev, N. M. Cohen, Q. Szabo, L. Fritsch, G. L. Papadopoulos, Y. Lubling, X. Xu, X. Lv, J.-P. Hugnot, A. Tanay, et al., Cell 171, 557 (2017).

[35] A. Javer, Z. Long, E. Nugent, M. Grisi, K. Siriwatwetchakul, K. D. Dorfman, P. Cicuta, and M. C. Lagomarsino, Nature Communications 4 (2013).

[36] A. Javer, N. J. Kuwada, Z. Long, V. G. Benza, K. D. Dorfman, P. A. Wiggins, P. Cicuta, and M. C. Lagomarsino, Nature Communications 5 (2014).

[37] E. Kepten, I. Bronshtein, and Y. Garini, Physical Review E 87, 052713 (2013).

[38] A. Amitai, A. Seeber, S. M. Gasser, and D. Holcman, Cell Reports 18, 1200 (2017).

[39] M. H. Hauer, A. Seeber, V. Singh, R. Thierry, R. Sack, A. Amitai, M. Kryzhanovska, J. Eglinger, D. Holcman, T. Owen-Hughes, et al., Nature Structural & Molecular Biology (2017).

[40] M. Barbieri, M. Chotalia, J. Fraser, L.-M. Lavitas, J. Dostie, A. Pombo, and M. Nicodemi, Proceedings of the National Academy of Sciences 109, 16173 (2012).

[41] A. S. Hansen, I. Pustova, C. Cattoglio, R. Tjian, and X. Darzacq, Elife 6, e25776 (2017).

[42] S. M. Gasser, Trends in Cell Biology (2016).

[43] A. Amitai, M. Toulouze, K. Dubrana, and D. Holcman, PLoS Computational Biology 11, e1004433 (2015).

[44] D. Hnisz, K. Shrinivas, R. A. Young, A. K. Chakraborty, and P. A. Sharp, Cell 169, 13 (2017).

[45] D. Peric-Hupkes, W. Meuleman, L. Pagie, S. W. Bruggeman, I. Solovei, W. Brugman, S. Gräf, P. Flicek, R. M. Kerkhoven, M. van Lohuizen, et al., Molecular cell 38, 603 (2010).

[46] A. Amitai and D. Holcman, Physics Reports 678, 1 (2017).

[47] S. C. Weber, A. J. Spakowitz, and J. A. Theriot, Proceedings of the National Academy of Sciences 109, 7338 (2012), http://www.pnas.org/content/109/19/7338.full.pdf.

[48] S. C. Weber, A. J. Spakowitz, and J. A. Theriot, Physical review letters 104, 238102 (2010).

[49] V. Dion and S. M. Gasser, Cell 152, 1355 (2013).

[50] A. Amitai and D. Holcman, bioRxiv, 076661 (2016).

[51] T. S. Harmon, A. S. Holehouse, M. K. Rosen, and R. V. Pappu, eLife 6 (2017).

## REFERENCES

[1] M. Bohn, D. W. Heermann, and R. van Driel, Physical Review E 76, 051805 (2007).

[2] J. Bryngelson and D. Thirumalai, Physical review letters 76, 542 (1996).

[3] E. P. Nora, B. R. Lajoie, E. G. Schulz, L. Giorgetti, I. Okamoto, N. Servant, T. Piolot, N. L. van Berkum, J. Meisig, J. Sedat, et al., Nature 485, 381 (2012).

[4] J. Dekker, K. Rippe, M. Dekker, and N. Kleckner, science 295, 1306 (2002).

[6] O. Shukron and D. Holcman, PLOS Computational Biology 13, e1005469 (2017).

[7] O. Shukron and D. Holcman, Physical Review E 96, 012503 (2017).

[8] M. Doi and S. Edwards, The Theory of Polymer Dynamics Clarendon (Oxford, 1986).

[9] A. Javer, Z. Long, E. Nugent, M. Grisi, K. Siriwatwetchakul, K. D. Dorfman, P. Cicuta, and M. C. Lagomarsino, Nature Communications 4 (2013).

[10] B. Bonev, N. M. Cohen, Q. Szabo, L. Fritsch, G. L. Papadopoulos, Y. Lubling, X. Xu, X. Lv, J.-P. Hugnot, A. Tanay, et al., Cell 171, 557 (2017).

[11] B. Eichinger, Macromolecules 13, 1 (1980).

[12] A. A. Gurtovenko and A. Blumen, in Polymer Analysis Polymer Theory (Springer, 2005) pp. 171–282.

[13] A. Amitai, A. Seeber, S. M. Gasser, and D. Holcman, Cell Reports 18, 1200 (2017).

[14] A. Javer, N. J. Kuwada, Z. Long, V. G. Benza, K. D. Dorfman, P. A. Wiggins, P. Cicuta, and M. C. Lagomarsino, Nature Communications 5 (2014).

[15] E. Kepten, I. Bronshtein, and Y. Garini, Physical Review E 87, 052713 (2013).

[16] N. Servant, N. Varoquaux, B. R. Lajoie, E. Viara, C.-J. Chen, J.-P. Vert, E. Heard, J. Dekker, and E. Barillot, Genome biology 16, 259 (2015).

[17] B. Langmead and S. L. Salzberg, Nature methods 9, 357 (2012).

[18] L. Giorgetti, R. Galupa, E. P. Nora, T. Piolot, F. Lam, J. Dekker, G. Tiana, and E. Heard, Cell 157, 950 (2014).

